# Life history scaling and the division of energy in forests

**DOI:** 10.1101/2020.06.22.163659

**Authors:** John M. Grady, Quentin D. Read, Sydne Record, Nadja Rüger, Phoebe L. Zarnetske, Anthony I. Dell, Stephen P. Hubbell, Sean T. Michaletz, Alexander Shenkin, Brian J. Enquist

**Affiliations:** National Great Rivers Research and Education Center, East Alton, IL, USA; Department of Integrative Biology, Michigan State University, East Lansing, MI, USA; Department of Biology, Bryn Mawr College, Bryn Mawr, PA, USA; National Socio-Economic Environmental Synthesis Center, Annapolis, MD, USA; German Centre for Integrative Biodiversity Research (iDiv) Halle-Jena-Leipzig, Deutscher Platz 5e, 04103 Leipzig, Germany; Department of Economics, University of Leipzig, Grimmaische Straße 12, 04109 Leipzig, Germany; Smithsonian Tropical Research Institute, Balboa, Ancón, Panama; Ecology, Evolutionary Biology and Behavior Program, Michigan State University, East Lansing, MI, USA; Washington University of St. Louis, Department of Biology, St. Louis, MO 63130, USA; Department of Ecology and Evolutionary Biology, University of California, Los Angeles, California, USA; Department of Ecology and Evolutionary Biology, University of Arizona, Tucson, AZ, USA; Earth and Environmental Sciences Division, Los Alamos National Laboratory, Los Alamos, New Mexico, USA; Department of Geography, Oxford University, Oxford, UK; The Santa Fe Institute, Santa Fe, New Mexico, USA

## Abstract

The competition for light has long been regarded as a key axis of niche partitioning that promotes forest diversity, but available evidence is contradictory. Despite strong tradeoffs between growth and survival with light, field tests suggest neutral forces govern tree composition across forest gaps and resource use across size classes. Here we integrate scaling and niche theory, and use data from >114,000 woody plants in a tropical, old growth forest to test and predict patterns of niche partitioning with size and light. Consistent with predictions, the relative abundance, production, light capture, and richness of species in life histories with fast growth follow a power law relationship, increasing 1–2 orders of magnitude along a solar and size gradient. Competitive neutrality between size classes emerges above the sapling layer, where increasing access to light is counterbalanced by stronger self-shading. Convergent power law patterns of resource partitioning across taxa and spatial scale suggest general life history tradeoffs drive the organization of diverse communities.

---

Competition for resources is regarded as key driver of forest assembly and diversity^1-5^, but empirical evidence is mixed. On the one hand, light declines to ∼2% intensity in hyperdiverse tropical forests^6^, and low light is associated with stunted growth and elevated mortality^7,8^. Niche theory posits that trait differences and associated tradeoffs promote niche partitioning and coexistence along resource gradients^5,9-11^. Indeed, in forests a *slow–fast* life history continuum has been observed in which light-demanding trees grow quickly in well-lit forest gaps (‘*fast’* life history), but at the cost of high mortality; conversely, *slow*-growing, long-lived trees are better able to recruit and survive in the dark understory^12,13^. Thus, *fast* trees may be competitively favored in high light areas, and *slow* species in the understory^14^, promoting coexistence. On the other hand, despite abundant experimental and demographic data documenting this tradeoff ^3,12,15^, field evidence for light-based niche partitioning with light is lacking, inspiring rival neutral theories^16^. For example, in tropical old-growth forests – where species richness is greatest – little variation was observed in the relative abundance and richness of pioneer and shade tolerant species across gap and non-gap sites^17,18^.

Indeed, recent work suggests that herbivory and pathogens, rather than resource competition, drive forest diversity^19-21^. Forest gaps increase light levels in the understory, but gaps are short-lived and stochastic. Further, shade-tolerant saplings are often already established when new gaps arise^22^, casting shade and blunting advantages un-germinated pioneers may hold. However, comparison of relative abundances of pioneer and shade tolerant species in understory gaps^17^ may bias results towards evaluation of saplings, which outnumber canopy trees by several orders of magnitude^23^. Although light along the forest floor varies unpredictably with gap occurrence, increasing light toward the canopy is a general feature of forests^6^. The most consistent gradient of light is vertical rather than horizontal, where larger individuals experience more consistent and prolonged light micro-environments. Over time, the sustained effects of light variation across size classes will exacerbate differences in growth and mortality rates between species. Thus, niche partitioning with light may only be apparent when ontogenetic size variation is explicitly modeled.

A scaling approach to forest structure quantifies the size dependence of abundance and resource capture from saplings to the canopy. Scaling relationships typically take a power law form, where *y ∝ M*^*α*^, *y* is an individual or population quantity, *M* is individual mass or diameter, and *α* is the slope on a log-log plot. Despite deviation in the smallest and largest organisms, global syntheses indicate that the population abundance of autotrophs – from phytoplankton to trees – declines with size at approximately the same magnitude that growth rate increases^24-26^. In particular, *R ∝ S*^*α*^, *A*_*i*_ *α S*_*i*_^*β*^, where *i* is size class, and α = –*β*. These opposing scaling relationships imply an ‘energy equivalence rule’ (EER), whereby individuals or species in different size classes collectively metabolize and grow at equal rates^24,27,28^. Because growth is fueled by assimilated resources, EER implies an additional dimension of competitive neutrality: across size classes, in addition to species. However, strong tests of EER are rare, as both growth and abundance scaling in local communities are seldom assessed in tandem.

Only recently has theory emerges to account for apparent size-neutrality in forests. The metabolic scaling theory of forests (MST)^29^ argues that EER emerges from space filling of tree crowns, in which each stem diameter size class has the same total leaf area, leading to equivalent rates of photosynthesis and respiration. Forest models incorporating more detail about canopy packing predict a similar size structure^2,30^, but even with equal leaf area per size class, EER appears paradoxical. Biomass production is a product of collective leaf area and resource availability, such as light. Larger trees receive more light on average^31^, and so should have higher production where light is limiting. Alternatively, niche partitioning may mitigate differences in light availability at upper and lower ends of the canopy, if high densities of shade tolerant species elevate understory production. However, with few exceptions^32^, scaling research has largely overlooked the role of resource availability and life history strategy needed to evaluate competition and niche partitioning. Even under EER size-neutrality, a vertical gradient of light availability may generate niche partitioning from the forest floor to the canopy.

Here we integrate niche and scaling theory to evaluate competition and niche partitioning in forests. We use recent competition theory^33^ to predict the division of resources and relative richness of species with opposing life history strategies. We test predictions using growth, abundance^34^, life history traits^35^, and light data^36^ for woody plants in a primary forest on Barro Colorado Island, Panama (BCI).

## A Scaling Model of Forest Assembly

To address the paradox of EER, we first note that competitive asymmetries with size are partially mitigated by the scaling of tree crown dimensions. Light first reaches the upper area of a tree crown, but declines as it penetrates the crown volume and is intercepted by leaves. Tree crowns are proportionally larger and deeper in taller trees, leading to more self-shading (Fig. S1). Thus, while the amount of light reaching tree crown tops is higher for taller trees, light interception per leaf will be more similar across size classes. Indeed, integrating a neighborhood model of light transmission^36^ with the scaling of LAI and crown depth^37^, we find that while average light reaching the crown declines 44–fold from the largest to smallest size class (Fig. 1a), the light per unit crown volume falls only by a factor of four (Fig. 1b, Table S1).

**Fig. 1.**
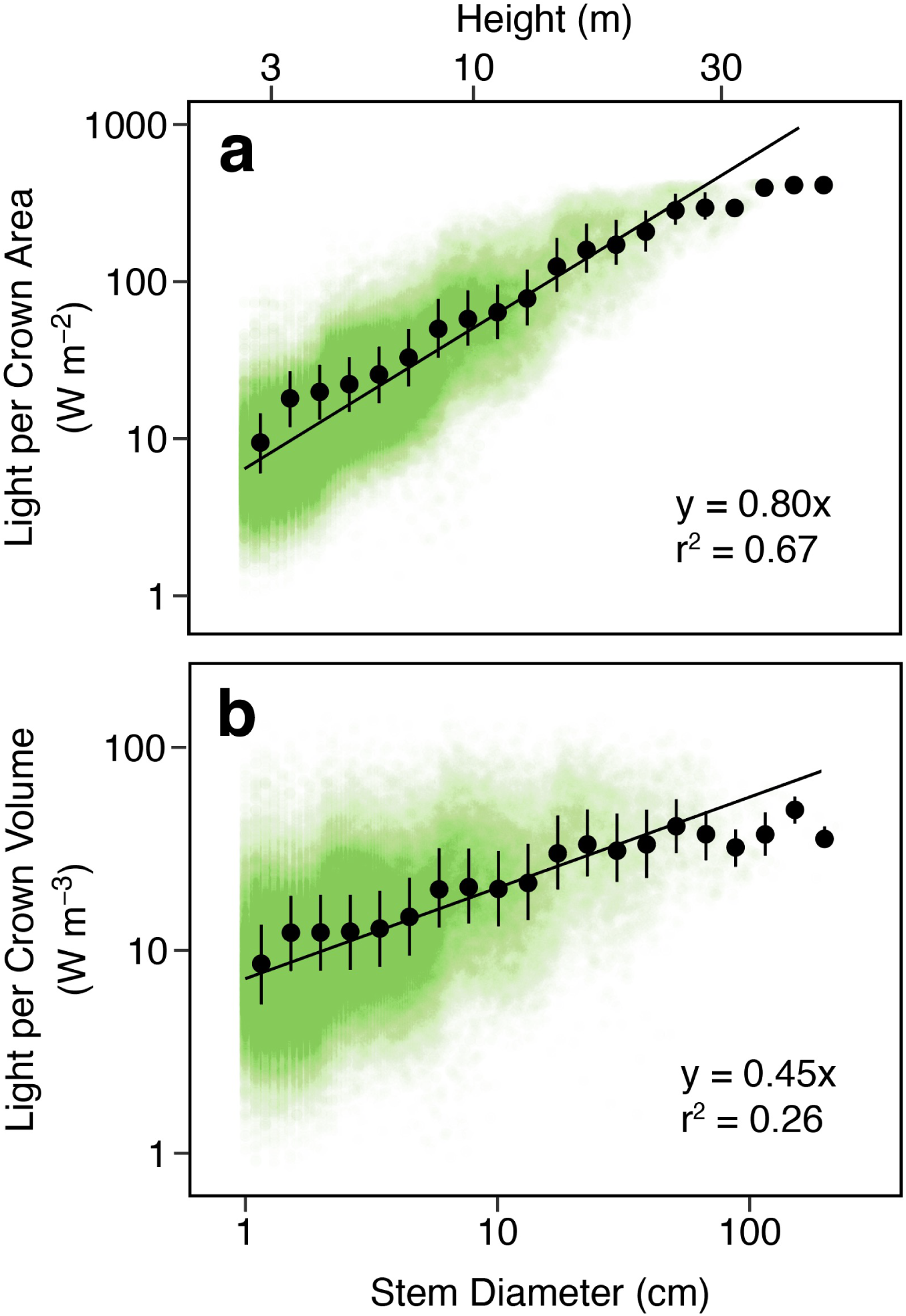
Light environment at crown and leaf. **a.** Individual light capture per crown area – a measure of light reaching a tree crown – increases over fortyfold from saplings to the canopy. **b**. LAI-weighted crown volume provides a better estimate of leaf level light and reveals a much more modest increase with stature: approximately fourfold. **a-b** mean values per size class are shown with 25% and 75% quantiles; n = 113,650. Data points are shown with 1% transparency for visualization; regressions with 95% credible bands are plotted. Height is shown as a reference, calculated from pooled-species scaling of stem diameter and height.

This remaining solar asymmetry with size, however, is predicted to drive niche partitioning. We use metabolic scaling theory (*MST*)^29^ as a baseline model to evaluate whole-community scaling slopes (Fig. 2, *gray*) and divergence by species in different life histories guilds (Fig. 2a). Treating growth as proportional to respiration, *MST* predicts individual growth *G* and abundance per area *A* to vary as a function of stem diameter *D* as:

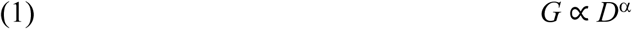

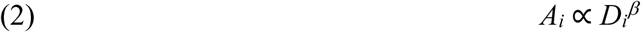

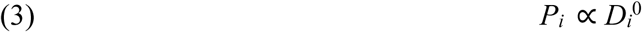

where α ∼ 2 and β ∼ –2, *i* is size class of stem diameter, *P* is total growth production in *i* (kg yr^−1^ ha^−1^ cm^−1^), and the slope is α + β. Because growth and metabolism are fueled by the uptake of limited resources, and variation in abundance drives concomitant resource use, we predict:

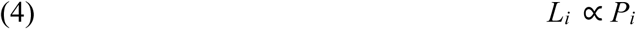

where *L*_*i*_ is intercepted light per size class (W m^−2^ cm^−1^), and *L*_*i*_ *α D*_*i*_^0^ where EER is observed. Deviation from power law scaling at the largest and smallest trees, however, is widespread ^24^ and may reflect constraints on maximum size, light limitation, and/or external mortality (e.g. canopy windthrow, drought^2^).

**Fig. 2.**
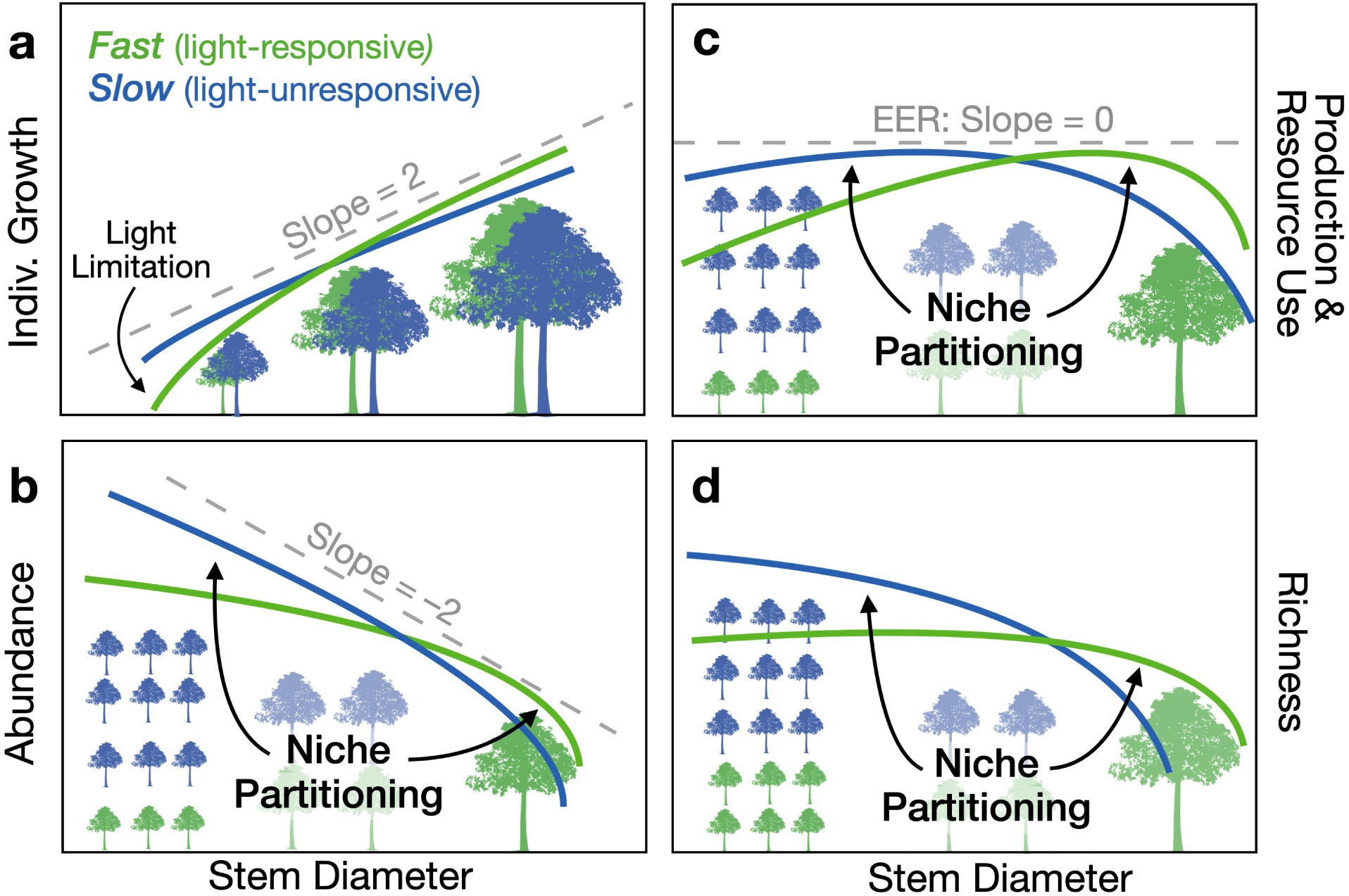
A scaling framework for life history variation and niche partitioning in forests. Light access covaries with tree size in forests, shaping patterns of niche partitioning across life histories. **a** The scaling of individual growth and resource uptake and **b** population abundance per area determines **c** aboveground production and the division of resources. **d** Richness is expected to tracking shifts in abundance (**b**) and resource capture **(c**). Scaling slopes for *all* individuals predicted by MST are shown in gray; life history guilds that vary in the responsiveness of growth and mortality to light are colored. Axes are log-transformed. Differences in scaling slopes between life histories imply niche partitioning, as *fast* species become relatively more abundant and speciose at larger sizes.

Life history tradeoffs in growth and mortality reflect, in part, linkages between metabolic performance and resource demand. *Fast* species have high leaf-level maximum assimilation that fuels rapid growth, but at the cost of elevated dark respiration^13,38^; a tradeoff that promotes energy deficit when light levels decline. This performance-resource tradeoff between *slow–fast* species implies systematic scaling deviations from Eqs. 1–3 in competing life history guilds. As light levels increase toward the canopy, elevated growth and declining mortality in *fast* species should lead to an increase in their relative abundance and production (Fig. 2a-c, *green*). Shade tolerant *slow* trees with high understory abundance (Fig. 2b, *blue*) may boost understory forest production and facilitate EER (Fig. 2c). As relative abundance and production shifts with size, we further predict accompanying shifts in relative richness (Fig. 2d). Competitive niche partitioning, or lack thereof, has been tested by examining whether relative abundance and richness of functional guilds change over resource gradients^17,18^. Thus, systematic shifts in relative abundance (Fig. 2b), relative production, resource uptake (Fig. 2c) and richness (Fig. 2d) with light would indicate resource-based niche partitioning between life history guilds.

More generally, species with a higher responsiveness to resource availability – steeper shifts in growth and mortality – are predicted to increase in relative abundance in high resource environments, and decline in low. This elevated growth–survivorship responsiveness is observed in a recently identified life history axis: a *recruitment–stature* life history (Fig. 3a, Fig. S2).

**Fig. 3.**
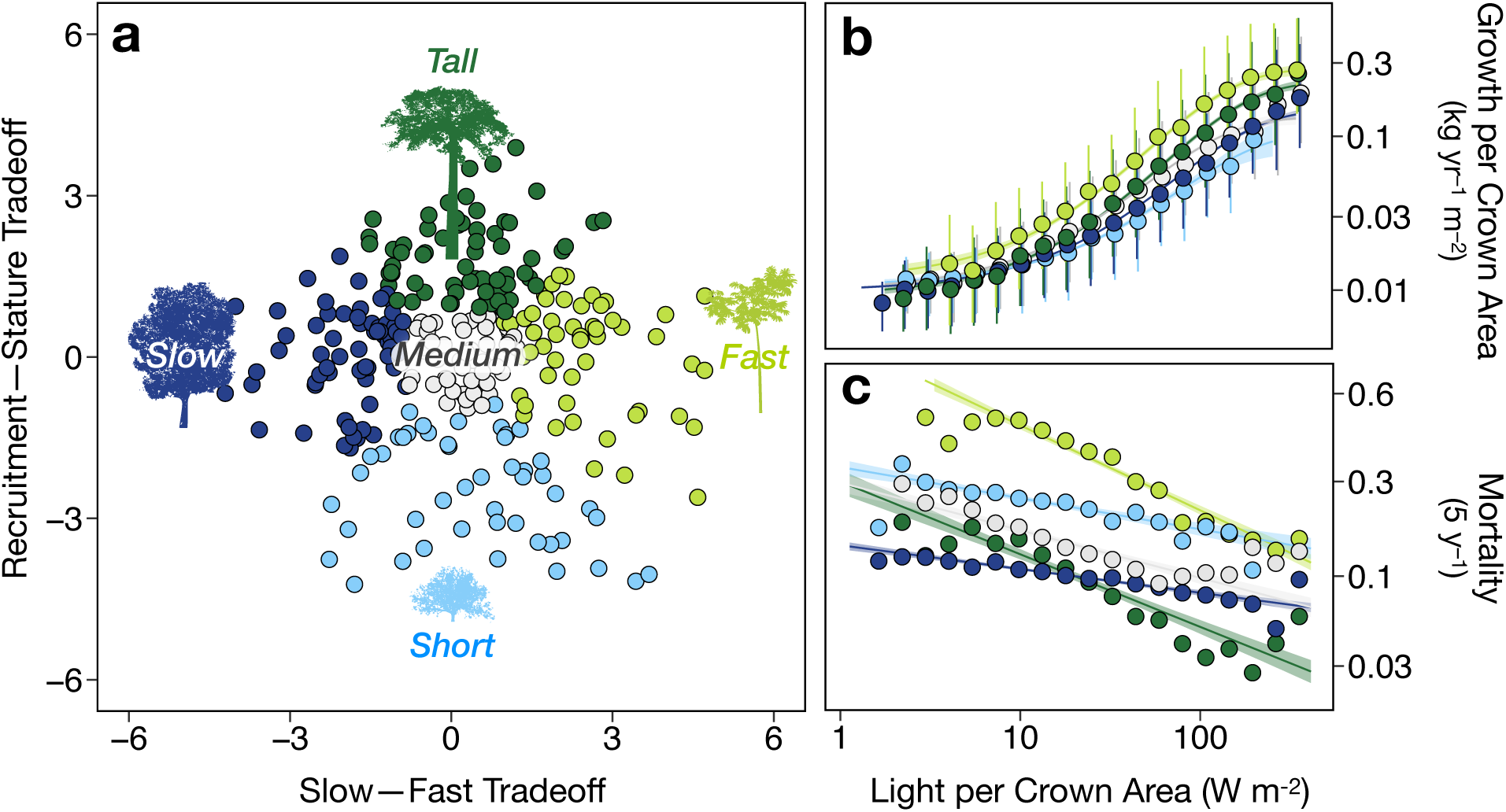
Life history guilds in woody plants. **a** Life history guilds in woody species at BCI are classified along two orthogonal axes: a *slow-fast* and *recruitment–stature* continuum. Each point represents a species at BCI in two-dimensional demographic space (PCA scores)^35^. Green life history guilds (*tall, fast*) are more responsive to high light environments, with steeper growth and mortality slopes in **b-c**. In **b**, Light availability is normalized for crown area; 25% and 75% quantiles shown for each binned value. For **b-c**, all points represent the geometric mean rate per size class, 95% credible bands are shown for all regression fits (n = 113,650).

Small, understory shrubs and trees (*short-lived breeders*, or *short*) have high recruitment rates, but small maximum size, slow growth, and high mortality^35^. Conversely, tall, *long-lived pioneers* (*tall*) grow quickly, reach a large stature and have long lifespans, but recruit poorly (Fig. 3a). The physiological traits characterizing *short* and *tall* species have received limited study, but despite differences in absolute mortality rates, *fast* and *tall* species share an elevated responsiveness to light (Fig. 3b-c, Figs. S3-S4, Table S2; mortality slope *fast* = –1.21 (Credible Interval: –1.29, – 1.12): mortality slope *tall* = –1.03 (CI: –1.15, –0.92); max. growth slope *fast* = 1.00 (CI: 0.97, 1.03), max. growth slope *tall* = 1.01 (CI: 0.99, 1.03)). Conversely, both *slow* and *short* species have shallower slopes and are comparatively unresponsive to increasing light (Fig. 3b-c, mortality slope *slow* = –0.34 (CI: –0.38, –0.29), mortality slope *short* = –0.44 (CI: –0.54, –0.35); growth slope *slow* = 0.88 (CI: 0.86, 0.89), growth slope *short* = 0.76 (CI: 0.68, 0.89)). This divergence in light responsiveness is consistent with the different light environments experienced by mature *short* and *tall* species, and imply similar scaling and niche partitioning shifts as *slow– fast* species in Fig. 2.

Recently, a metabolic competition model was advanced to explain a global power law in relative richness and the division of resources between animal competitors with fast (*F)* and slow (*S)* metabolism^33^. The proportion of assimilated resources (*Y*) shifted from *F to S* along a resource gradient (*r*), where *Y*_*F*_*/Y*_*S*_ *α r*^*σ*^. Extending this model to trees and substituting from Eq. 5, we predict the dimensionless ratio of production between life history guilds follows a power law with light availability:

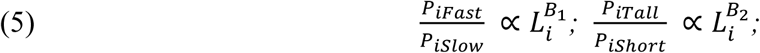

where *B*_*1*_ and *B*_*2*_ are scaling exponents > 0. Because light availability follows a power law with size (Fig. 1), we expect a similar power law shift across diameter as well. Following ^33^, we predict concurrent shifts in relative richness *R* across a light gradient, but at shallower rates, reflecting the sublinear relationships between abundance and richness^39^:

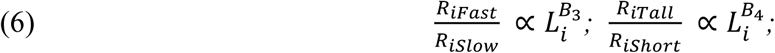

where *B*_*1*_ > *B*_*3*_, and *B*_*2*_ > *B*_*4*_

Thus, we make the following predictions:

*H1: Divergent scaling across life histories*. Approximate MST slopes (Eqs. 1-3) and EER occurs for shade tolerant *slow* and all woody trees collectively (*all*) at intermediate sizes, but divergent scaling growth, abundance, production and richness slopes, including deviation from EER, will occur in light-responsive *fast* and *tall* species, consistent with niche partitioning (Fig. 2a-d).

*H2: Resource-production symmetry.* Linking metabolism to resource use, collective light capture rates with size will match total production (Eq. 5, Fig. 2c), with matching scaling slopes of production and light for respective life history guilds.

*H3: Power law niche partitioning.* The ratio of abundance, production and richness by opposing life histories will follow a power law with stem diameter and light (Eq. 5, Eq. 6).

We use piecewise, regression fits to quantify scaling shifts with size, and evaluate scaling patterns against stem diameter at breast height (dbh; 1.3 m). We focus on scaling patterns at intermediate size (∼ 3 – 50 cm dbh), which comprise the majority of the vertical range of tree heights (∼5 – 25 m height). Population values for abundance, production and average growth rate are logarithmically binned for visualization, following White *et al*^40^.

## Scaling Patterns

Growth, abundance and production slopes are near MST predictions (Eqs. 1–3) for all individuals within the forest (*all*), but diverge across life history guilds (Fig. 4a-c, Table S3). As predicted (*H1*), *fast* and *tall* species have significantly steeper abundance, total production slopes than *short, slow* and *all* species (Fig. 4b-c, Fig 2, Table S4), and richness is relatively higher at larger size classes (Fig 4d). However, scaling shifts in individual growth rate with size were not observed, and can be described with a single power law (Fig. 4a vs Fig. 4b, Table S4, Fig. S5). This may reflect tradeoffs in leaf max. assimilation rate with leaf number in shade tolerant vs intolerant species^38^. Indeed, significant differences in biomass growth with life history are only apparent when evaluated across light intensities (Fig. 3b). Across the whole community (*all*), production scaling at intermediate sizes is near but somewhat higher than energy equivalence (slope = 0.18, CI: 0.16–0.19, Fig. 4c), with marked declines in the smallest and largest sizes. This value falls within the range observed by meta-analyses that support EER as a central tendency across sites^24,27^ (see also values across sampling periods in Fig. S6a). As predicted, only shade tolerant species approach EER, although a modest decline in production at larger sizes coincides with an increasing role by *fast* and *tall* species (Fig. 4c, Table S4). Representing 65% of all individuals, shade tolerant *slow* species largely drive scaling patterns of abundance of *all* trees collectively, and are closest to MST predictions among life history strategies (Fig. 4).

**Fig. 4.**
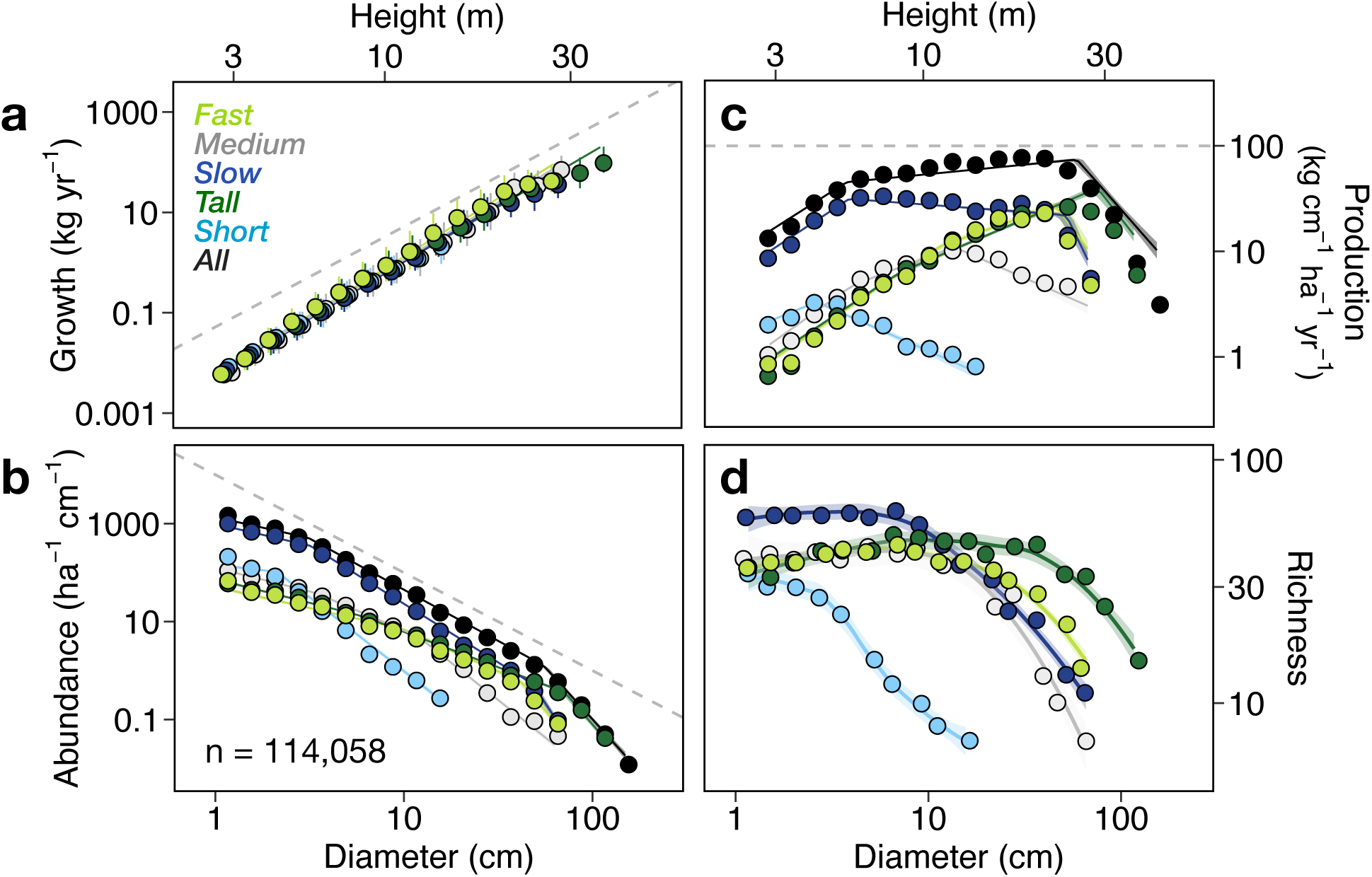
Scaling of growth, density, production and richness across life history guilds. Plots of observed scaling relationships with dbh, with the *MST* predicted slope indicated by dashed gray lines. In **a**, each data point is the mean value per size increment, with 25% and 75% quantile bars. In **b** and **c**, plotted values are summed values per size class, with three-part piecewise regression fits. **a-c**, 95% credible bands are shown for all regressions.

## Size Neutrality & Niche Partitioning with Light

Production scaling does not differ significantly from the scaling of light capture (Fig. 4c vs Fig. 5a, Fig. 5b), supporting our symmetry hypothesis (*H2*) and indicating that life history variation in production scaling reflects equivalent variation in resource use. Thus, convergence toward competitive neutrality in light capture occurs between size classes for *all* individuals from ∼3 – 50 cm stem diameter, or ∼5 – 25 m height (Table S5, Fig. 5, Fig. S7). Corresponding to light capture, the scaling of total crown volume approaches equivalence, but is somewhat lower, with a modest decline at intermediate sizes (Fig. S6b, Table S6). An observed twofold decline in total crown volume for *all* individuals partially offsets the fourfold increase in light availability with size (Fig. 1b), pushing the system toward EER.

**Fig. 5.**
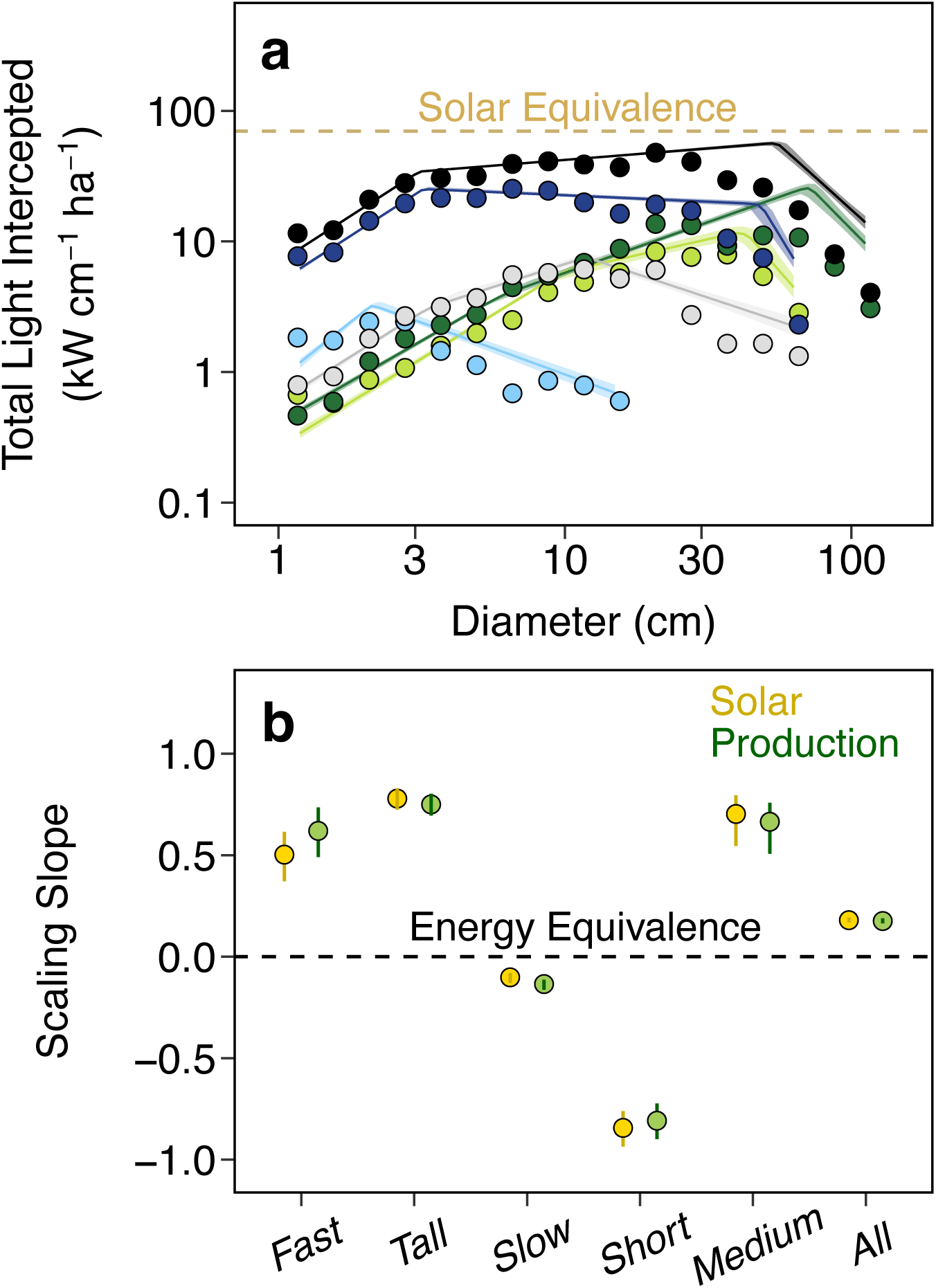
Production and resource uptake symmetry. **a** The scaling of light capture across life history guilds with 95% credible bands. **b** Light capture slopes at intermediate sizes from **a** is similar to the scaling slope of production in Fig. 4c for respective life history guilds, as indicated by overlapping 95% credible intervals.

Shifts in absolute and relative abundance, production and light capture are consistent with competitive tradeoffs and niche partitioning (*H3*; Fig. 4 vs Fig. 2). Mean life history PCA scores shifts monotonically toward *faster* and *taller* values and in larger size classes and higher light intensity (Fig. S8). *Slow* and *short* species dominate light capture at small sizes, and light-responsive *fast* and *tall* life histories gaining an increasing share of light resources toward the canopy (Fig. 4c, Fig. 5a). Indeed, despite considerable curvature in abundance and production scaling in small and large individuals (Fig. 4b-c), shifts in relative abundance, production, and richness between *fast* vs *slow* and *tall* vs *short* are nearly linear in log space, supporting *H3* (Fig. 6a-b, Table S7-S8). This is particularly striking for relative production. Production for every life history guild plunges nearly two orders of magnitude at the lowest light levels (Fig. S9c) even as relative share remains approximately linear (Fig. 6a), following Eq. 5. Absolute richness is curvilinear with diameter (Fig. 4d), but plotting the ratio of richness yields a continuous power relationship (Fig. 6 c-d), as light-responsive guilds become at least equally speciose in the canopy. As predicted, the slopes of relative richness is directionally similar to abundance and production, but with shallower slopes (Fig. 6 c-d vs Fig. 6a-b, Table S8).

**Fig. 6.**
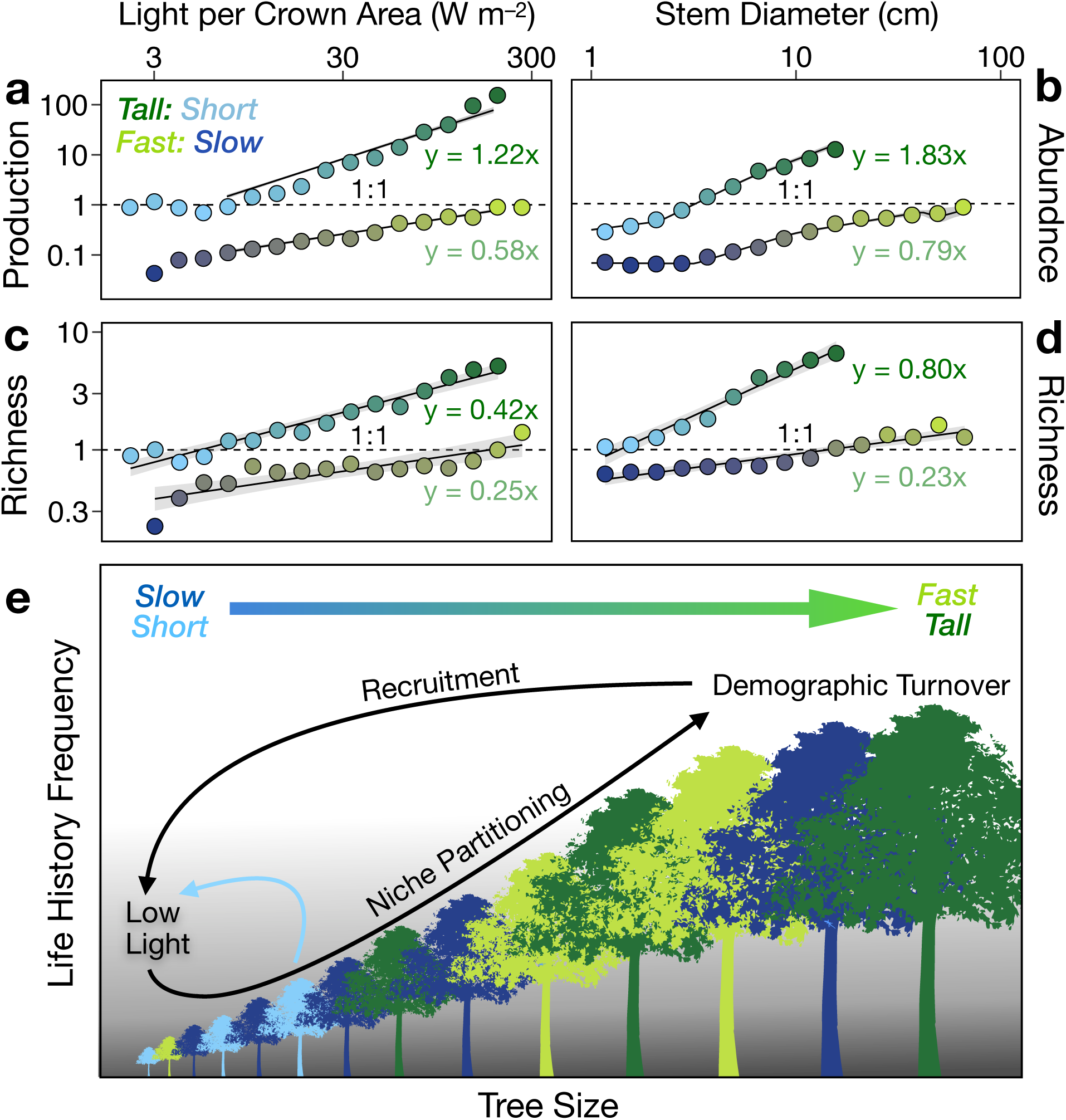
Life history scaling with size and light. Light availability increases with tree size, driving life history shifts in *tall* and *fast* **a** relative production, **b** relative abundance and **c-d** relative richness. Each point represents the ratio of values for opposing life histories at a size class; the dashed line signifies a ratio of 1. Slopes and 95% credible bands are shown for **a-d**. For **a**, abundance scaling with light is nonlinear below 7 W m^−2^, constraining the regression fit. In **b**, the slope at intermediate sizes slope for piecewise regression is shown. **e** Schematic representation of niche partitioning based on life history scaling. Larger *fast* and *tall* individuals outcompete *slow* and *short* individuals toward the well-lit canopy, leading to demographic turnover. Max. size constraints also limits the size-frequency of *short* understory species (blue arrow). Low survival or recruitment of *fast* and *tall* species in the dimmer understory resets relative abundance back to *short* and *slow*, implying a perpetual cycling of life history frequency with size.

The observed demographic shift from light-responsive to light-unresponsive species has parallels to succession^41^, but over vertical space rather than time. Our analyses focuses on an old-growth plot with no recorded history of stand clearing or significant disturbance^42^. Demographic analyses indicate the abundance of pioneer and climax guilds is near equilibrium, with annual disturbance rate of 0.43% to 1% of area^43^. The continuous shifts in relative abundance and resource capture with size are consistent with sustained niche partitioning rather than succession. Despite competitive advantages realized by *fast* and *tall* trees toward the canopy, these species are poor recruiters (*tall*) or suffer high mortality in the understory (*fast*), driving the relative abundance of saplings shifts back toward *slow* and *short* species along the forest floor. A cycling of life history frequency with size promotes coexistence and may typify closed forests (Fig. 6e).

Our results help resolve the paradox of forest energy equivalence and the role of light in community assembly. First, the large crown volumes of canopy trees increase self-shading, reducing the availability of light to leaves of larger individuals even in well-lit portions of the forest, and limiting the competitive advantages of a larger size (Fig. 1a-b). Second external mortality in the canopy creates gaps that promote light penetration, sapling recruitment^2^, and vertical ‘space-filling’ of tree crowns that reduce crown volume and light asymmetries with size (Fig. S6b). Finally, niche partitioning across a light gradient plays a role in maintaining high production from the understory to the canopy. Except at the smallest sizes, *slow* and understory *short* species capture sufficient energy in the lower canopy to push the system toward EER (Fig. 4c). *Fast* and *pioneer* forms deviate from EER but increase total energy capture for all trees in the upper canopy. Indeed, despite representing only 15% of individuals, *fast* and *tall* species together produce as much annual biomass as *slow* species that are over four times more abundant.

## Synthesis

Metabolic scaling approaches have attracted interest for revealing law-like patterns in metabolism and abundance that link cells to ecosystems^44,45^. However, such scaling relationships have been argued to hold limited relevance to community ecology and coexistence theories, because at the scale of communities the role of stochastic processes, resources, and interspecific variation are thought to be more important^46^. Conversely, community ecologists often overlook how trait frequency and resources shift with organismal size.

Integrating approaches can reveal general rules for community organization. Tradeoffs in size and metabolism with abundance – as exemplified by EER – are ubiquitous, shaping community structure across the globe^47^. Less well known is how variation in metabolism shapes species interactions and community structure. We show approximate power law relationship of richness and niche partitioning in competing plants at a local scale that mirrors global patterns observed in animals^33^. In both cases, fast-growing/fast-metabolizing species systematically increase their share of available resources and relative richness in higher resource environments. Faster metabolism corresponds with faster biological work rates (e.g., growth, movement, reproduction)^48^ that offer competitive advantages if sufficient resources can be procured^13,33,49^. Although competition has long been a central focus of ecological theory, moving beyond species-pair evaluations is a challenge^50^. Emergent scaling shifts between competing guilds offer a path forward, highlighting shared mechanisms that underly the assembly of diverse communities.

## Acknowledgements

We thank K.C. Cushman for sharing stem allometry data, and H. Muller-Landau and C.E. Farrior for discussions.

## Author Contributions

JMG conceived the study, JMG developed theory, JMG and QDR wrote the paper, JMG, QDR, NR, and BJE designed the approach, QDR and JMG performed analyses, NR and SPH contributed data. All authors discussed results and edited the manuscript.

## Code and data availability

All R and Stan scripts required to reproduce our analysis is available at https://github.com/qdread/forestscalingworkflow, and an R package containing additional functions is available at https://github.com/qdread/forestscaling. BCI survey data are publicly available through the Smithsonian Institution (https://repository.si.edu/handle/10088/20925). For demographic data used to determine life history guilds, refer to Rüger et al. 2018. All other data we used were taken from the literature and are cited in the methods.

## List of Supplementary Materials

### Materials and Methods

Pdf document. Includes method description, an embedded summary table of major scaling patterns (Table S3) and 10 figures.

**Supplementary References.** References 48 – 71.

**Supplementary Table 1. Fitted parameters for light capture per crown area and volume by diameter.** Parameters correspond to Fig. 1. q025 and q975 represent 2.5% and 97.5% quantiles of parameter estimates, corresponding the 95% credible intervals.

**Supplementary Table 2. Scaling parameter summary and credible intervals for growth rate by light and life history.** Parameters corresponds to Fig. 3. r^2^ values are also provided.

**Supplementary Table 3. Slopes with credible intervals for major scaling patterns.** Embedded in Materials and Methods document.

**Supplementary Table 4. Scaling parameter summary for growth, abundance, and production.** Parameters correspond to Fig. 4. Widely Applicable Information Criterion (WAIC) also provided.

**Supplementary Table 5. Scaling parameter summary and credible intervals for individual and total light interception by diameter and life history.** See also Fig. 5a, S7.

**Supplementary Table 6. Scaling parameter summary and credible intervals for total crown volume by diameter and life history.** Parameters correspond to Fig. S6b.

**Supplementary Table 7. Scaling parameter summary and credible intervals for growth, abundance, and production by light and life history.** Parameters correspond to Fig. S9.

**Supplementary Table 8. Production and abundance ratios.** Parameters correspond to Fig 6a-b.

## Materials and Methods

### Site and Demographic Data

We used long-term demographic data from a mesic neotropical forest on Barro Colorado Island (BCI), Panama (9°9’N, 79°51’W). BCI is warm throughout the year (average daytime and nighttime temperatures of 32°C and 23°C, respectively), with most of the 2500 mm of rainfall falling during a wet season, from April to November^51^. Censuses of all free-standing woody stems ≥ 1 cm dbh, (diameter at breast height, measured 1.3 m from ground), at 0.1 cm resolution, have been conducted on a 50 ha portion of the island at five year intervals since 1980; see ^52^ for full description.

We focus on a primary forest without history of disturbance. For this reason, we excluded approximately 2 ha of the survey area consisting of secondary forest^53^. In addition, individuals within 20 m of the plot edge were excluded, as their light environment could not be fully described because not all of their neighbors within a 20 m radius were mapped^36^. All analyses are conducted on the 42.84 ha remaining of the 50 ha survey area after excluding the secondary forest and the area within 20 m of the plot edge. We show abundance from 1995, growth from 1990 to 1995, and light estimates from 1995. Patterns were qualitatively are similar to other census years (see Fig. S6a). Both abundance and growth analyses were based on the 114,058 trees tagged in both 1990 and 1995. The 132,982 trees, which includes the new 1995 recruits, were used for plotting total production (binned values in Fig. 4c). We imputed the values for the ∼18,000 new 1995 recruits that were not present in 1990, using the growth by diameter scaling regressions. Total growth regression was calculated from abundance and individual growth regressions.

### Processing and Forest Survey Data

Our quality control procedure follows ref ^54^. First, we removed 6 strangler fig species (< 100 individuals total) because their growth form prevents accurate estimation of biomass from diameter. Tree ferns and woody lianas were not included in the census protocol. We corrected the diameter of 11 species that have tapering or buttressed stems, whose diameters were measured at a higher point than breast height (1.3 m). For buttress correction, we used species-specific parameters provided by K. C. Cushman (personal communication) that were fitted using models described in references ^55-57^. Specifically, we used the taper parameter *b* for each species and applied the following correction to estimate the corrected diameter from the observed diameter: *d*_*corrected*_ = *d*_*measured*_*e*^*b*(*h*−1.3)^, where *h* is the height at which diameter was measured.

### Allometry from Tree Diameter Measurements

We estimated the following measurements for all individual trees using allometric functions of dbh: tree height, crown area, crown depth, crown volume, and aboveground biomass.

We calculated diameter-height allometry and diameter-crown area allometry using parameter values generated from measurements taken on BCI^58^. The generalized Michaelis-Menten function was found to be the best allometry for tree height, as it reaches an asymptote at high dbh values: 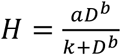, where *D* is dbh and *a, b*, and *k* are constants. For crown area, a power law relationship fit best: *A* = *aD* ^*b*^, where *D* is dbh and *a* and *b* are constants. In both cases we used species-specific coefficients provided from ref. ^58^ where possible, and the all-species coefficients for species not included in the study (9.1% of individuals).

To calculate diameter-crown depth allometry, we obtained data from the authors of ref. ^59^ and used them to fit a power law relationship to each species. For the species without individual measurements, we calculated an allometry for each of the five life history guilds and used the group-specific allometry (191 species 30.1% of individuals). For species without either individual measurements or life history assignment, we used the all-species allometry (14 species representing 1.1% of individuals).

Allometries for height, crown area, and crown depth were corrected for Jensen’s inequality to eliminate biases resulting from log transformation (see “Estimating Total Growth Scaling Relationships” below). We took the correction factors for tree height and crown area from ^58^. We calculated the crown depth correction factors for each life history guild, using the formula from ^60^.

We estimated crown volume from crown diameter and depth, assuming a half-elliptical crown shape for all individual trees, following ref ^59^: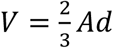, where *A* is crown area and *d* is crown depth.

Next, we estimated aboveground biomass of all individuals given dbh and height. The parameters of the allometry are taken from ref (Rüger, Comita, Condit, Purves, Rosenbaum, Visser, Joseph Wright, et al. 2018)^61^ and were developed to apply across moist tropical forests: *AGB* = 0.0509*GD*^2^*H*, where *G* is wood specific gravity, *D* is dbh, and *H* is tree height. This approach uses wood specific gravity measurements at the genus or species level (measurements provided by K.C. Cushman, personal communication).

We took the difference in aboveground biomass allometries between consecutive censuses to estimate 5-year biomass growth for each tree for each census. We converted this to annual biomass growth rate using the following equation: 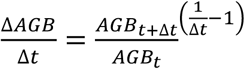 where Δ*t* is the census interval in days (roughly 5 years). We excluded individuals appearing for the first time in the tree census from the individual growth rate analyses (14% of all individuals), although these individuals were included in all other analyses. We removed outliers where trees were recorded as gaining more than 20 cm dbh between two censuses or were entered into the census with > 10 cm dbh following the first census, representing likely errors. This resulted in removing < 300 individuals of 114,058.

### Binned Data

Population abundance and production per size class are not individual properties but collective ones. Thus, for empirical and visual representation we plot binned values for each size class. All regression fits, however, are based on raw data. In Fig. 4a, individual growth rates are binned and plotted for comparison to collective properties, and because life history differences are not visually distinguishable when plotting > 114,000 individual data points. Where binned individual values are plotted, 25% and 75% quantile bars are also plotted to show variation around the mean. We also provide heat maps of individual rates in the supplements (Figs. S8, S12). To bin data, we follow White et al. ^40^ by plotting summed abundance and total growth that is measured over plant size bin increments in logarithmic space and then divided by the bin range to show arithmetic mean densities and growth rates per cm diameter. Thus, binned values of production represent the arithmetic mean of total growth per unit stem diameter across a logarithmic range. All plotted points have a minimum of 20 individuals per bin.

### Life History Classification

We use life history data from Rüger et al. 2018 (ref ^35^), following their classification scheme of *fast, medium, slow, short-lived breeder (short)*, and *long-lived pioneer* (*tall*). Rüger *et al.* analyzed 282 species at BCI, using demographic data across four canopy layers. They observed a *fast*−*slow* continuum, or growth−survival tradeoff that corresponded to previous assessments of shade tolerance^12^, where shade-tolerant species have slow life histories (slow maturation, long lifespan) and shade-intolerant species have fast life histories (fast stem diameter growth, short lifespan; Fig. S2). Rüger *et al* also analyzed per capita recruitment rates and found evidence of a second stature−recruitment tradeoff, in which slow-growing shrubs and small trees had short lifespans, but high recruitment rates. Conversely, long-lived pioneers had low recruitment rates, but grew fast and survived well, and thus attained large adult stature. A *medium* strategy was observed within these extremes. Using data from ref ^35^, species scores in weighted PCA, including growth, survival, and recruitment rates were used to classify life histories (Fig. 3a). Specifically, both PCA axes were normalized by the absolute value of the 10^th^ and 90^th^ percentile values on that axis divided by 2. All species within a radius of 1 from the origin were included in the *medium* group. The remaining species were divided into quadrants at a 45° angle from the PCA axes, resulting in five life history guilds.

### Quantifying Individual Light Capture

To characterize the light environment for individual trees, we utilized published light estimates derived from annual censuses of vegetation density (http://richardcondit.org/data/canopy/bciCanopyReport.php). The presence/absence of vegetation was measured along a 5 m grid across the 50 ha plot and at six height intervals: 0-2, 2-5, 5-10, 10-20, 20-30, and ≥ 30 m. If vegetation was present, it was assumed to cast shade in the same manner as a 5 m flat circle at the vertical midpoint of each height range. A shade index was calculated from the proportion of the sky obscured by vegetation accounting for its angle and distance from a focal tree. The proportion of open sky reaching a given tree was converted to the proportion of light reaching a tree by linking the shade index to irradiance measurements taken at many locations on BCI but outside the study plot (see ^62^, and for a full description of the algorithm see ^36^). The proportion of irradiance reaching each tree was multiplied by average overhead insolation at the latitude of BCI (418 W m^−2^ at 9°9’ N) following ref ^63^ to obtain the incoming light energy per area reaching the vertical projection area of the crown of each tree. 408 individuals lacked a modeled light value, or 0.36%.

To convert light energy per unit area to total incoming light energy reaching each tree’s crown, we multiplied incoming light energy per unit area by crown area. To convert total incoming light energy to light energy per unit crown volume, we divided incoming light energy by crown volume and then accounted for self-shading within the tree canopy using the Beer-Lambert equation: *p* = 1 − *e*^−*KLAI*^. Here, *p* is the proportion of light penetrating a tree canopy, *k* is the light extinction coefficient, and LAI is the leaf area index. We use *k* = 0.5 from Kitajima *et al* ^37^ taken from four Panamanian trees, and calculated as the slope of light transmittance against LAI (Fig. S1b). One atypical species – pioneer *Cecropia longipes* – was excluded. *Cecropia* accumulates virtually no LAI or crown depth as it grows^37^. For our analysis, LAI scaling with crown depth was applied to species-specific data on crown depth to estimate individual LAI at a given size. We derived the parameters of this scaling relationship (*LAI* = 0.368*D*^0.367^, where D is our allometrically derived measurement of crown depth, Fig. S1a) from the measurements provided for the four Panamanian tree species in Kitajima *et al* ^37^. Parameters were estimated using hierarchical mixed-effects regression to log LAI vs log crown depth, with species identity as a random intercept.

### Relationship between Mortality Rate and Light

The BCI census protocol records whether individual trees died between censuses. We used the tree status codes from the BCI censuses in 1990 and 1995 to fit logistic regressions to individual tree mortality as a function of incoming light per unit crown area. We fit a hierarchical logistic regression in a Bayesian framework, fitting random slopes and random intercepts for each functional guild, and using a logit link function. The model was specified as follows:

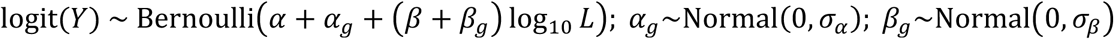

Where *Y* is a binary outcome variable equal to 1 if the tree died during the five-year period, *L* is the light received per unit crown area in units of W m^−2^, *g* is an index representing the functional guild of the individual, α is the global intercept (fixed effect), α_j_ is the guild-specific intercept (random effect), β is the global slope (fixed effect), and β_j_ is the guild-specific slope (random effect). We set normal priors on the fixed and random effects, and exponential priors on the standard deviations of the effects.

To visualize the mortality data, we binned the set of all individuals across the 1990 and 1995 censuses in equally sized intervals in logarithmic space and calculated the fraction of individuals in each functional guild that died between the two surveys.

### Scaling Analysis: Individual Growth

To quantify individual growth scaling we used a single power law (Eq.1), and also considered a two-segment, hinged power law to assess potential deviation (see A. Gelman, https://statmodeling.stat.columbia.edu/2017/05/19/continuous-hinge-function-bayesian-modeling/). The functional forms of the power law and two-segment hinged power law follow, along with a description of the parameters of each function:

∘ 1-segment power law:
  ▪ log *G* = log *β*_0_ + *β*_1_ log *D* + *ε*; *ε*∼*Normal*(0, *σ*)
  ▪ Can also be expressed as 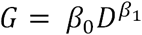
  ▪ Parameters (3):
    - *β*_0_ intercept
    - *β*_1_ slope
    - *σ* standard deviation of error distribution
∘ 2-segment hinged power law:
  ▪ 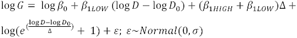
  ▪ Parameters (6):
    - *β*_0_ intercept
    - *β*_1*LOW*_ slope at lower values of D
    - *β*_1*HIGH*_ slope at higher values of D
    - *D*_0_ cutoff between low and high slopes
    - Δ smoothing parameter for hinge
    - *σ* standard deviation of error distribution

We used the following prior distributions on the parameters of the individual growth functions:

- 1-segment power law
  ∘ *β*_0_: Lognormal(1, 1)
  ∘ *β*_1_: Lognormal(1, 1)
  ∘ *σ*: Exponential(0.1)
- 2-segment hinged power law
  ∘ *β*_0_: Lognormal(1, 1)
  ∘ *β*_1*LOW*_: Lognormal(1, 1)
  ∘ *β*_1*HIGH*_: Lognormal(1, 1)
  ∘ Δ: Exponential(10)
  ∘ *D*_0_: Lognormal(1, 1)
  ∘ *σ*: Exponential(0.1)

A simple power law was best supported (Table S4), and results are based on a single power law fit unless stated otherwise.

### Scaling Analysis: Abundance

We fit three functions to the abundance distributions: a Pareto or single-segment power law, a two-segment piecewise power law that captures the steeper downward slope for large trees, and a three-segment piecewise power law that allows three different slopes for small, midsize, and large trees. For the Pareto distribution, we fixed the *D*_*min*_ parameter to 1 cm, the minimum diameter recorded in the tree censuses, where 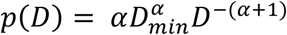 (note that when the slope is −2, *α* = 1). We used the functional form of the two-segment power law or double Pareto from the literature^64^. We modified the two-segment power law further by adding an additional segment at the lower tail to obtain the three-segment power law. The functional forms of the power law, two-segment power law, and three-segment power law follow, along with a description of the parameters of each function and constants calculated as a function of the parameters.

∘ 1-segment power law, also known as Pareto distribution:
  ▪ 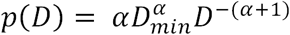
  ▪ Parameters (1):
    - −(*α* + 1) slope
  ▪ Constants (1):
    - *D*_*min*_fixed at 1 cm
∘ 2-segment piecewise power law, also known as double Pareto distribution:
  ▪ 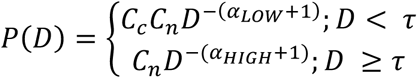
  ▪ Parameters (3):
    - −(*α*_*LOW*_ + 1) slope at lower values of D
    - −(*α*_*HIGH*_ + 1) slope at higher values of D
    - *τ* cutoff between low and high slopes
  ▪ Constants (3):
    - 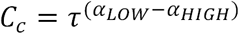 (continuity constant to ensure that the function is continuous at *τ*
    - 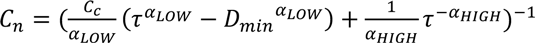 normalization constant to ensure that the function has an integral equal to 1
    - *D*_*min*_fixed at 1 cm
∘ 3-segment power law:
  ▪ 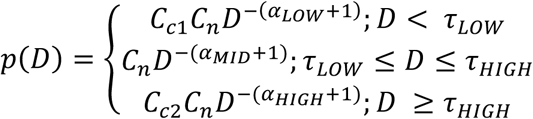
  ▪ Parameters (5):
    - −(*α*_*LOW*_ + 1) slope at lower values of D
    - −(*α*_*MID*_ + 1) slope at middle values of D, between the two cutoffs
    - −(*α*_*HIGH*_ + 1) slope at higher values of D
    - *τ*_*LOW*_ cutoff between low and middle slopes
    - *τ*_*HIGH*_ cutoff between middle and high slopes
  ▪ Constants (4):
    - 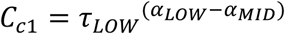 continuity constant to ensure that the function is continuous at *τ*_*LOW*_
    - 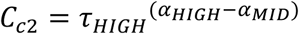 continuity constant to ensure that the function is continuous at *τ*_*HIGH*_
    - 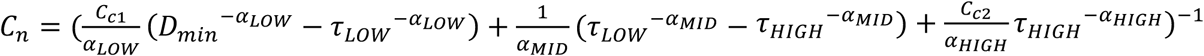 normalization constant to ensure that the function has an integral equal to 1
    - *D*_*min*_fixed at 1 cm

We used the following prior distributions on the parameters of the abundance functions:

- Abundance: 1-segment power law
  ∘ *α*: Lognormal(1, 1) truncated between [0, 5]
- Abundance: 2-segment power law
  ∘ *α*_*LOW*_: Lognormal(1, 1) truncated between [0, 10]
  ∘ *α*_*HIGH*_: Lognormal(1, 1) truncated between [0, 5]
  ∘ *τ*: Uniform truncated between minimum and maximum diameter values observed
- Abundance: 3-segment power law
  ∘ *α*_*LOW*_: Lognormal(1, 1) truncated between [0, 10]
  ∘ *α*_*MID*_: Lognormal(1, 1) truncated between [0, 5]
  ∘ *α*_*HIGH*_: Lognormal(1, 1) truncated between [0, 5]
  ∘ *τ*_*LOW*_: Uniform truncated between minimum and maximum diameter values observed
  ∘ *τ*_*HIGH*_: Uniform truncated between *τ*_*LOW*_ and the maximum diameter value observed, ensuring that *τ*_*HIGH*_> *τ*_*LOW*_

### Abundance Scaling Relationships

We fit one model for each of the three functional forms described above. Each of the three models were fit for all trees together (*all*), as well as for each of the five life history guilds separately and for the individuals not classified in a group, in the 1995 census. We compared the three models within each group using the Widely Applicable Information Criterion^65^ (WAIC; Table S4), which is asymptotically equivalent to Bayesian leave-one-out cross-validation. We calculated point estimates and 95% credible intervals for all model parameters, as well as the point estimates and 95% credible intervals for the fitted values of abundance. All fitted values were computed at 101 diameter values evenly spaced in log-space. For visual representation, we converted the abundance functions into units of individuals per hectare by multiplying the fitted values by the total number of individuals in the appropriate life history guild divided by 42.84 ha (study area with edge and secondary forest excluded; see above).

### Individual and Total Growth Scaling Relationships

We also evaluated individual growth scaling patterns using hierarchical Bayesian models. We fit the two functional forms described above to all trees together, each of the five life history guilds, and the individuals not classified in a group, all using the values from the 1995 census. We compared the fits using WAIC (Table S4). We calculated point estimates and 95% credible intervals for all model parameters, as well as the point estimates and 95% credible intervals for the fitted values of individual growth, at the same diameter values as we did for abundance. In addition, we calculated the Bayesian R-squared value for the individual growth fits ^65^.

### Total Growth Production Scaling Relationships

At each sample from the posterior distribution, we took the product of the fitted values for abundance and individual growth at a range of sizes to calculate the total growth curve for each combination of abundance and individual growth fits, yielding six combinations (1, 2, or 3 segment power law for abundance × 1 or 2 segment power law for individual growth). For the figures where the total growth fits are visualized, we removed the bias introduced by the log-transformation used in the individual growth regression. The bias is a result of Jensen’s inequality ^66^ (where total growth in a size class is consistently underestimated by assuming all individuals in the size class have the mean size); it affects only the intercept of the total growth function. We calculated the correction factor (from ^60^) as follows. First, we found the standard error of the estimate *SEE* from the log-log regression:

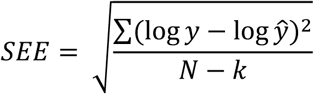

Here, *y* are the observed values, *ŷ* are the fitted values (linear predictor), *N* is the number of values, and *k* is the number of parameters (for example, 2 in the case of the log-log regression). The correction factor *CF* is a function of the standard error of the estimate:

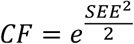

We calculated the correction factor for each Monte Carlo iteration and multiplied the fitted values by the median value of *CF* across the Monte Carlo samples to correct the visualization. This correction did not affect our statistical inference regarding the individual or total growth slopes, because it only influences the intercept of the individual growth function.

We also calculated the fitted values and 95% credible intervals of the slopes of the abundance, individual growth, and total growth functions by numerically calculating the slopes between each of the 101 points at which we evaluated the functions. The uncertainty around the fitted values for the total growth scaling includes uncertainty from both the size-growth relationship and the size-abundance relationship.

### Total Light and Total Crown Volume Scaling Relationships

We also estimated the scaling relationships between total light energy received and diameter, and between total crown volume and diameter. We used the same methods as for total growth scaling: we regressed individual light energy received on diameter using the one-segment and two-segment power law functions as we did for individual growth. We multiplied the fitted values of individual light received and abundance together to yield the total light scaling relationship. We calculated the same model statistics for individual and total light as we did for individual and total growth. The same procedure was followed for total crown volume.

### Relationship between Normalized Light and Normalized Growth

For light-growth curves we used a resource response function where growth rate increases sigmoidally with size until reaching a maximum rate, following a von Bertalanffy formulation:

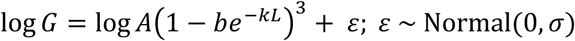

where *G* is growth per unit crown area (kg yr^−1^ m^−2^) and *L* is light received per unit crown area (W m^−2^), accounting for partial shading of the crown by other trees. *A, b*, and *k* are parameters. Because light capture will reflect crown area as well as light intensity, we normalized for crown area differences by dividing both growth and light capture by crown area in Fig. 3b-c. As above, we fit the function in a Bayesian framework.

To find maximum rate of increase of *G* relative to *L*, we calculate the slope at the inflection point of the function G = *A*(1 − *be*^−*KL*^)^3^ on a log-log plot, i.e. the maximum of 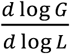.

Using the identity 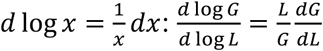. We find

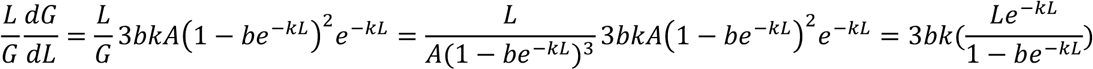

To find the maximum value of 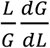, we differentiate the above function and find the value of *L* where the derivative is equal to zero 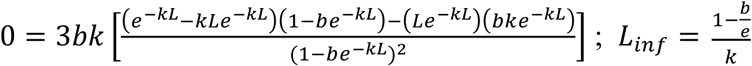 where *L*_*inf*_ is the value of *L* at the inflection point. We evaluate the function G = *A*(1 − *be*^−*KL*^)^3^ at that value to get *G*_*inf*_, the value of *G* at the inflection point, where 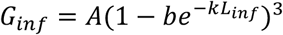 We evaluate 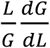 at *L*_*inf*_ to get the value of the maximum slope of the log-log plot.

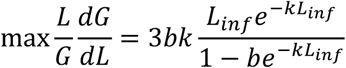

### Relationship between Normalized Light and Size

We fit log-linear regressions to the relationship between incoming light per unit crown area and diameter, and to the relationship between incoming light per unit crown volume and diameter. As above, the models were fit in a Bayesian framework.

### Abundance and Growth Scaling Relationships, Scaled by Light per Crown Area

We also fit abundance and growth scaling relationships where, instead of using diameter to represent individual size, we used incoming light per unit crown area to quantify the scaling behavior of abundance, individual growth, and total growth as light increases. For abundance scaling, we fit a power law similar to the 1-segment abundance power law described above, but we fixed the *D*_*min*_ parameter at 7 W m^−2^, roughly where power law behavior begins (see Fig. S9). For growth scaling, we fit a log-linear regression identically to the 1-segment growth model described above. The model fitting procedure was the same as for the other fits described above, and we used the same procedure to calculate total growth scaling as the product of individual growth and abundance scaling, which corrects for bias introduced by Jensen’s inequality.

### Richness scaling

Richness is an emergent property that does not reflect individual rates or cumulative values like abundance. Thus, richness in Fig 4d was calculated summing unique species occurrence in the plot for a log binned size class, with loess regression fits. Ratios of these values were plotted in Fig. 6c-d; linear regression fits were performed on binned data.

### Light and Production Scaling

At the leaf level, assimilation saturates at some light intensity. However, approximate proportionality of light interception and growth in Eq. 4 is predicted to occur for the following reasons. First, most trees are not in the upper canopy where light is highest, so production will generally increase with higher light. Second, even in the upper canopy, many leaves are below the uppermost crown and experience self-shading. Thus, at the whole tree level, greater light will increase assimilation rates in leaves lower in crown and thereby increase tree growth. Indeed, individual growth rate does not plateau at highest observed light intensities, but slows only modestly (Fig. S3). Finally, both total light capture and production per size class are proportional to the number of individuals in that class, facilitating a proportional relationship to each other.

### Theoretical Predictions

Previously, we showed that fast-metabolizing, endothermic mammals and birds have performance advantages over slow-metabolizing shark and bony fish competitors ^33^. In particular, water temperature mediates the availability of prey resources: cold water slows the metabolism and speed of fish prey, leading to easier capture of fish by marine endotherms. From the tropics to the poles the relative abundance of mammals and their consumption of prey follows a power law with water temperature, with the observed slope matching model predictions. Further, the ratio of endotherm richness to ectotherm richness follows globaly follows a directionally similar but shallower power law (slope of 0.79 vs 1.05). More generally, elevated metabolism corresponds with a variety of faster ‘work’ rates – e.g., growth and development rate ^67,68^, reproduction ^69,70^, and locomotion ^33^ – but at the cost of higher resource demand. Generalizing across taxa, performance-demand tradeoffs are predicted to lead to power law shifts between competing life history guilds in forests over a resource gradient, whereby species with ‘acquisitive’ strategies and high leaf metabolism ^13,71^ increase in relative abundance and resource uptake in high light environments.

### Model Fitting Details

All models were coded in the Stan language and fit with CmdStan 2.15.0. The model outputs were visualized and the model statistics were calculated using R 3.6.0, including the rstan package version 2.18.2 (ref ^72^) and the loo package version 2.1.0 (ref ^73^) for finding information criteria. In all cases, we used Hamiltonian Monte Carlo to sample from the posterior distribution, with 3 chains, 5000 warmup samples per chain which we discarded, and 1000 post-warmup samples per chain which we retained. We assessed convergence of posterior distributions by visually examining trace plots and by ensuring that 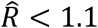 for all parameters ^74^.

## Supplementary Table

**Table S3.** Scaling exponents (slopes) for growth, light capture and abundance.

| **Life History** | **Growth** | **Indiv. Light** | **Abundance** | **Production** | **Solar Capture** | **Spp** | **n** |
| --- | --- | --- | --- | --- | --- | --- | --- |
| *Fast* | 2.33<br>(2.30,<br>2.35) | 2.21<br>(2.20,<br>2.22) | -1.71<br>(-1.84,<br>-1.60) | 0.620<br>(0.491,<br>0.735) | 0.504<br>(0.371,<br>0.615) | 52 | 8,898 |
| *Tall* | 2.20<br>(2.18,<br>2.21) | 2.22<br>(2.21,<br>2.24) | -1.45<br>(-1.50,<br>-1.40) | 0.751<br>(0.695,<br>0.803) | 0.780<br>(0.724,<br>0.830) | 65 | 11,136 |
| *Slow* | 2.26<br>(2.26,<br>2.27) | 2.30<br>(2.29,<br>2.30) | -2.39<br>(-2.42,<br>-2.38) | -0.134<br>(-0.163,<br>-0.113) | -0.101<br>(-0.130,<br>0.081) | 64 | 86,864 |
| *Short* | 2.25<br>(2.22,<br>2.28) | 2.22<br>(2.19,<br>2.24) | -3.02<br>(-3.10,<br>-2.94) | -0.807<br>(-0.899,<br>-0.724) | -0.843<br>(-0.936,<br>-0.761) | 44 | 10,456 |
| *Medium* | 2.29<br>(2.27,<br>2.31) | 2.33<br>(2.32,<br>2.35) | -1.63<br>(-1.79,<br>-1.54) | 0.665<br>(0.507,<br>0.758) | 0.704<br>(0.545,<br>0.795) | 52 | 12,817 |
| *All* | 2.26<br>(2.26,<br>2.26) | 2.27<br>(1.26,<br>2.27) | -2.09<br>(-2.10,<br>-2.08) | 0.176<br>(0.164,<br>0.189) | 0.181<br>(0.168,<br>0.192) | 293 | 132,982 |

2.5% and 97.5% quantiles (i.e., 95% credible intervals) are shown in parentheses. Abundance, production and solar capture slopes are for the intermediate portion of three-part, piecewise Pareto fits, ~ 3 – 50 cm dbh. Light slopes represent the increase in light interception with stem diameter. Light capture and production refer to collective rates across stem diameters. 2,809 individuals (2.1% of total) were unclassified and are included in analysis of *All*. n represents the number of individuals over two sampling periods (1990, 1995) used to calculate total growth production ('Production'). All slopes are from log-log regression fits.

## Supplementary Plots

**Fig. S1.**
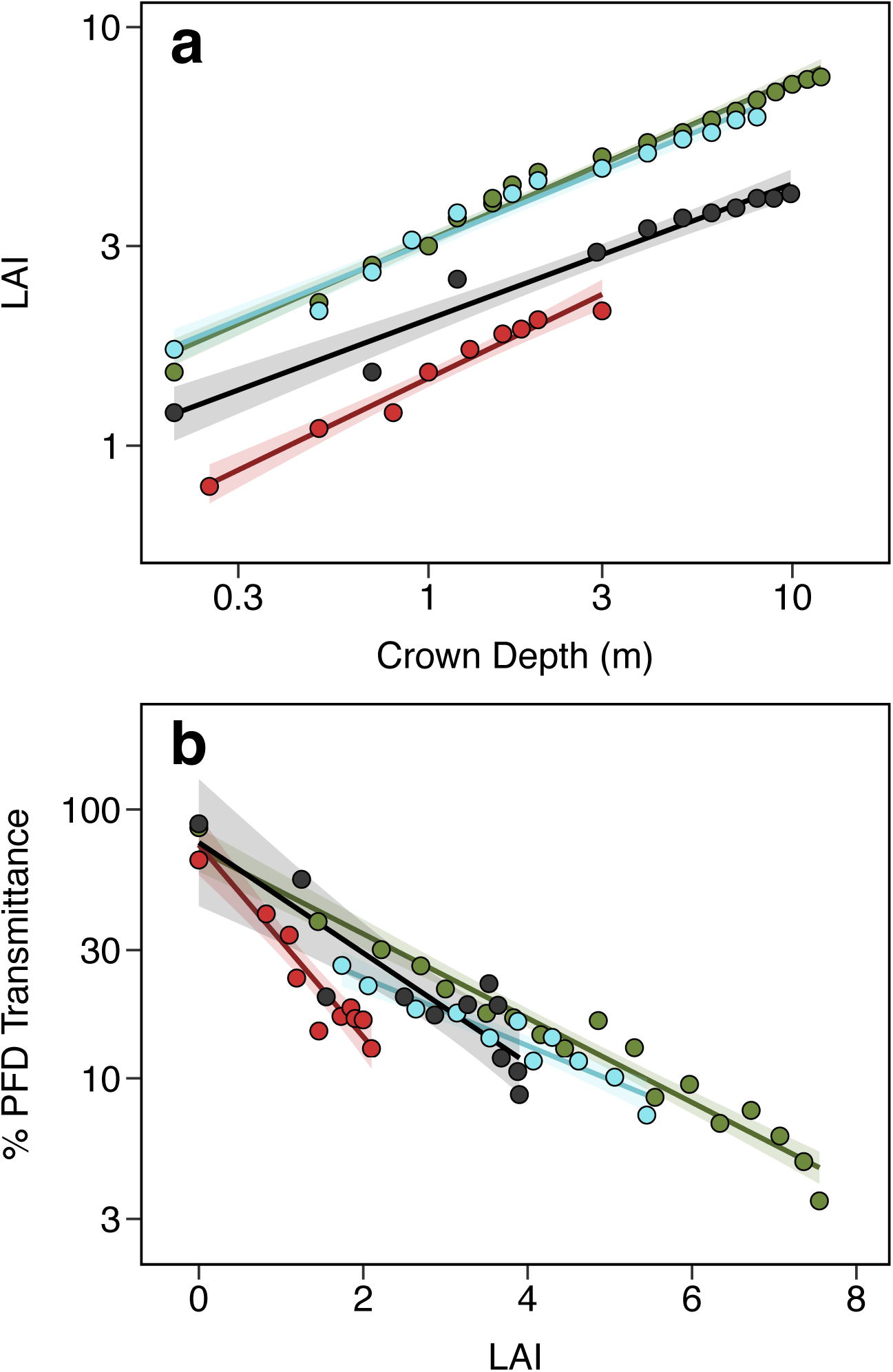
Light transmittance with tree size. **a** Leaf area index (LAI) increases with crown depth. **b**. High LAI lead to declines in photosynthetic flux density (PFD). Data for four Panamanian trees, from Kitajima et al (2005). Full species names are *Anacardium excelsum, Luehea seemannii, Antirrhoea tricantha*, and *Castilla elastica*.

**Fig. S2.**
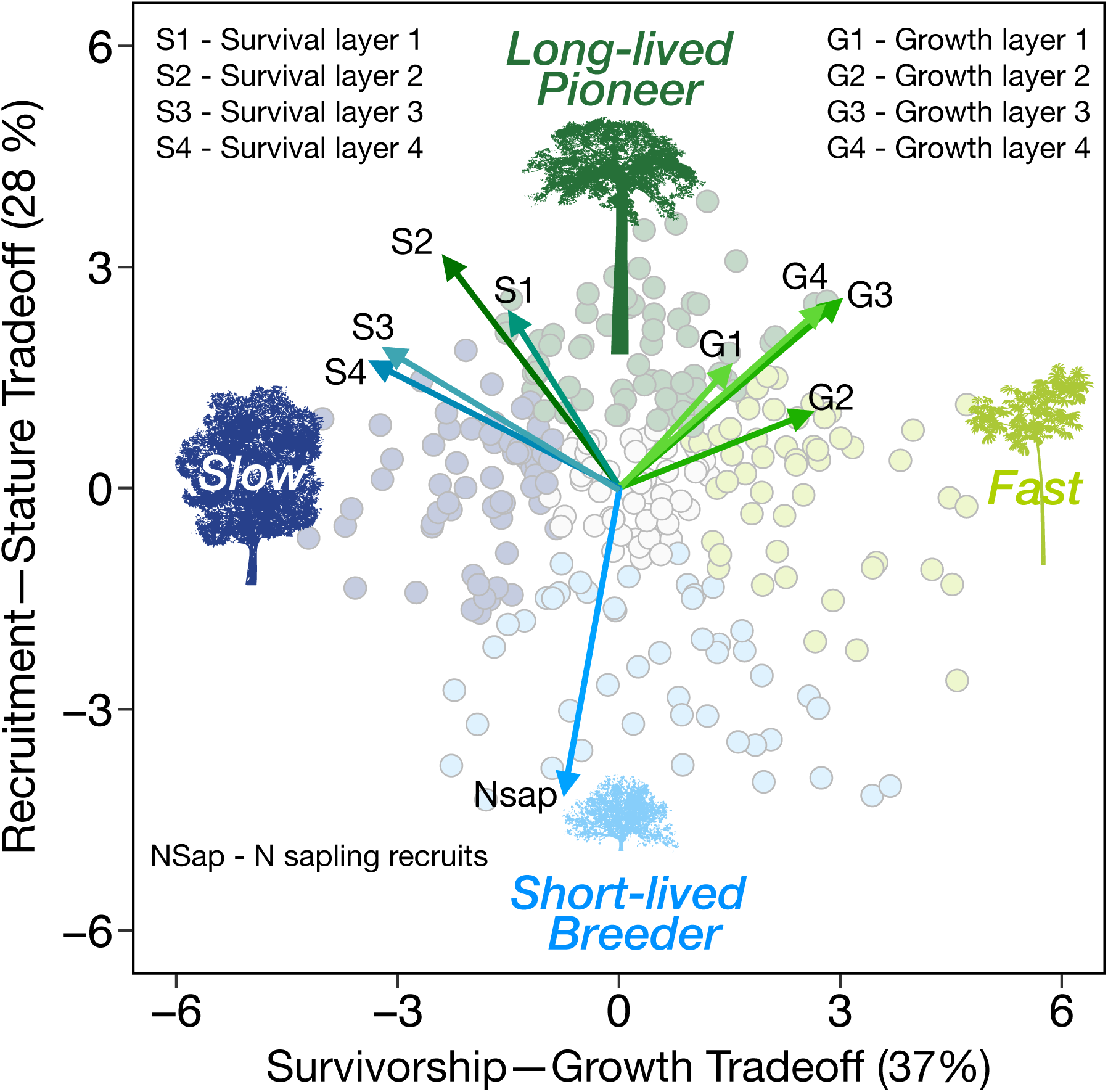
Life history Principal Component Analysis (PCA) scores with loadings. Data from Ruger *et al*, 2018, correspond to their Fig. 1a. Survival and growth loadings are shown at varying canopy layers, where higher layer numbers reflect higher heights. Growth is change in stem diameter per basal area per time. Classification protocol for life history guilds are described in Methods.

**Fig. S3.**
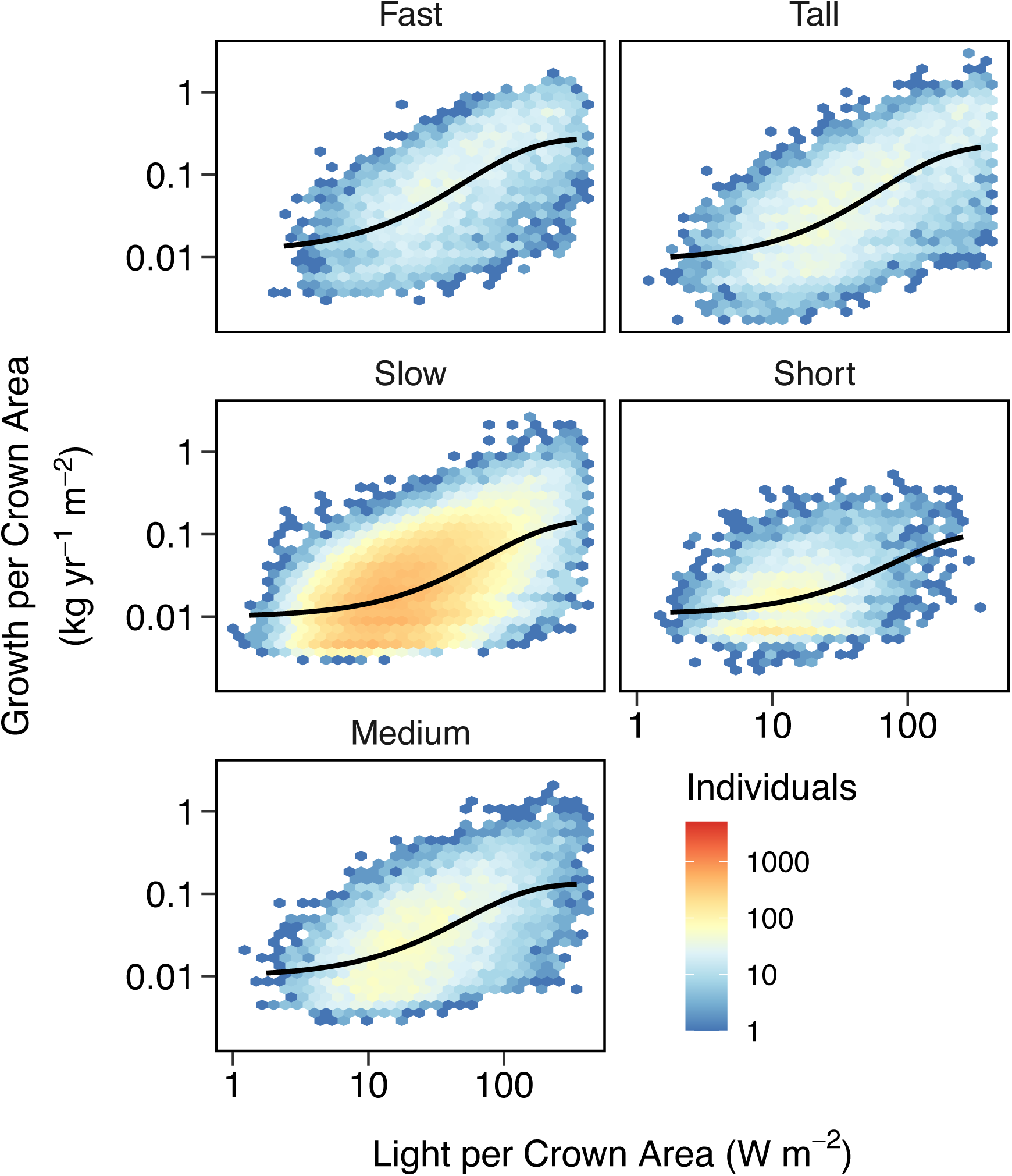
Heat map of growth per light capture. Higher light intensity leads to more rapid growth that slowly saturates. Tree size effects on growth rate and light capture are normalized by dividing by tree crown area. Number of individuals per cell are indicated by color.

**Fig. S4.**
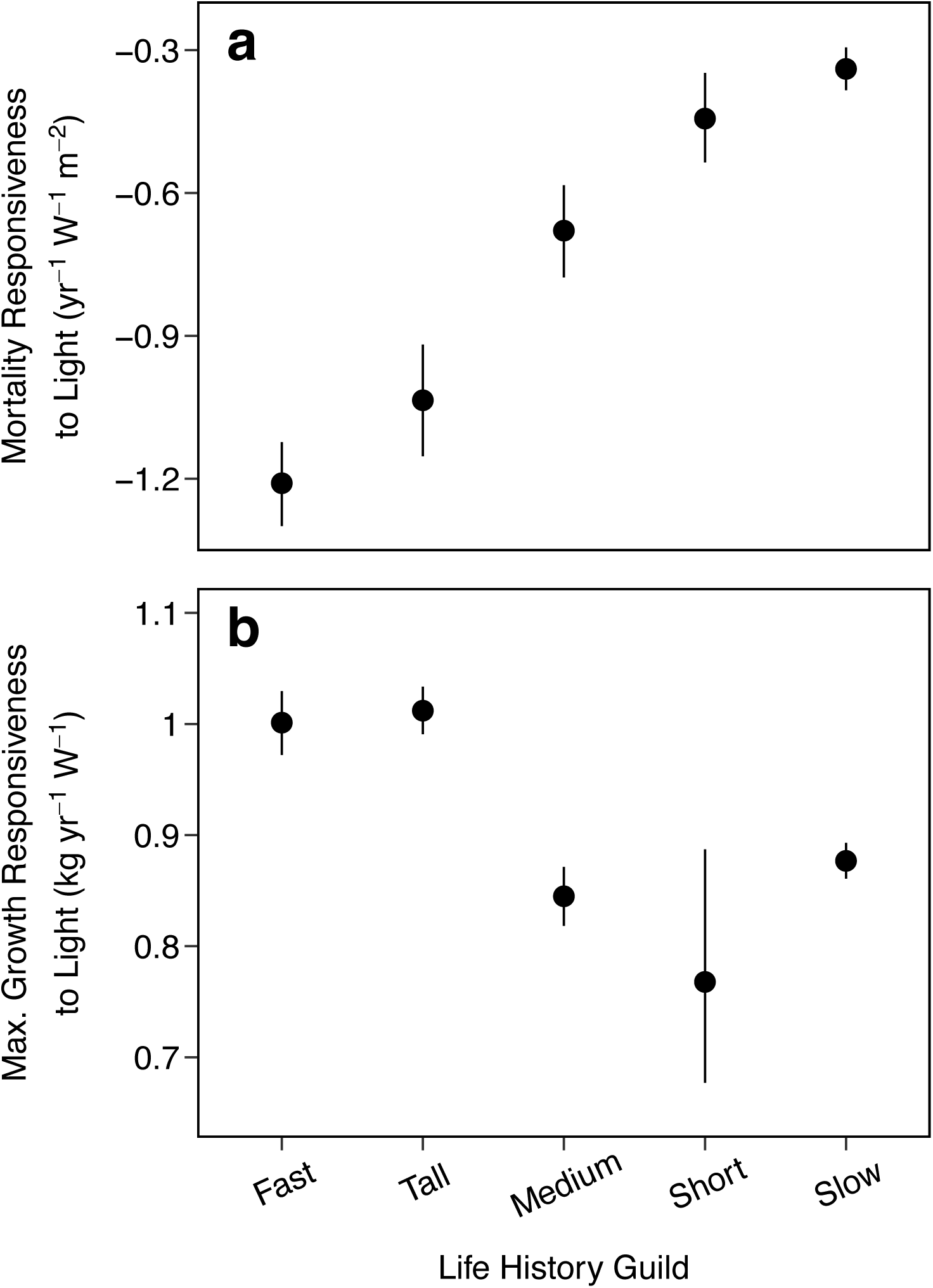
Growth and mortality responsiveness to light. Life history guilds vary in how their mortality (**a**) and growth rate (**b**) changes with light intensity (Fig. 2a-b, Fig. S5). *Fast* and *tall* species are the most responsive (steeper slopes): mortality declines rapidly in high light, while growth increases rapidly in high light. **b** Shown are the steepest increases in growth rate with light in Fig. S5. **a-b** Error bars indicate 95% credible intervals.

**Fig. S5.**
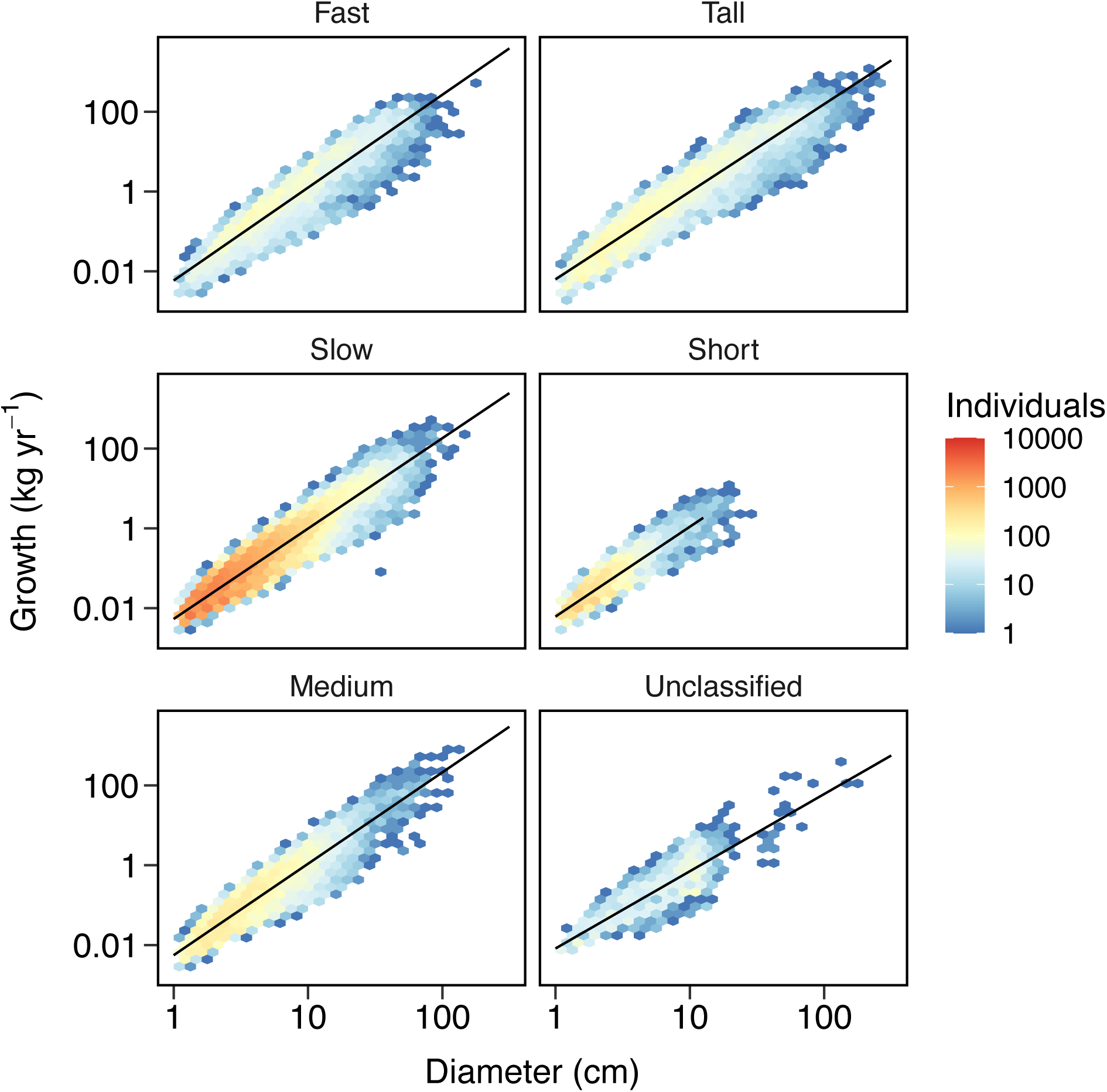
Heat map of individual growth scaling per life history guild. The density of individuals for a given size and growth rate is shown. A reference slope (dashed) of 2 predicted by MST is shown. See also Table S4.

**Fig. S6.**
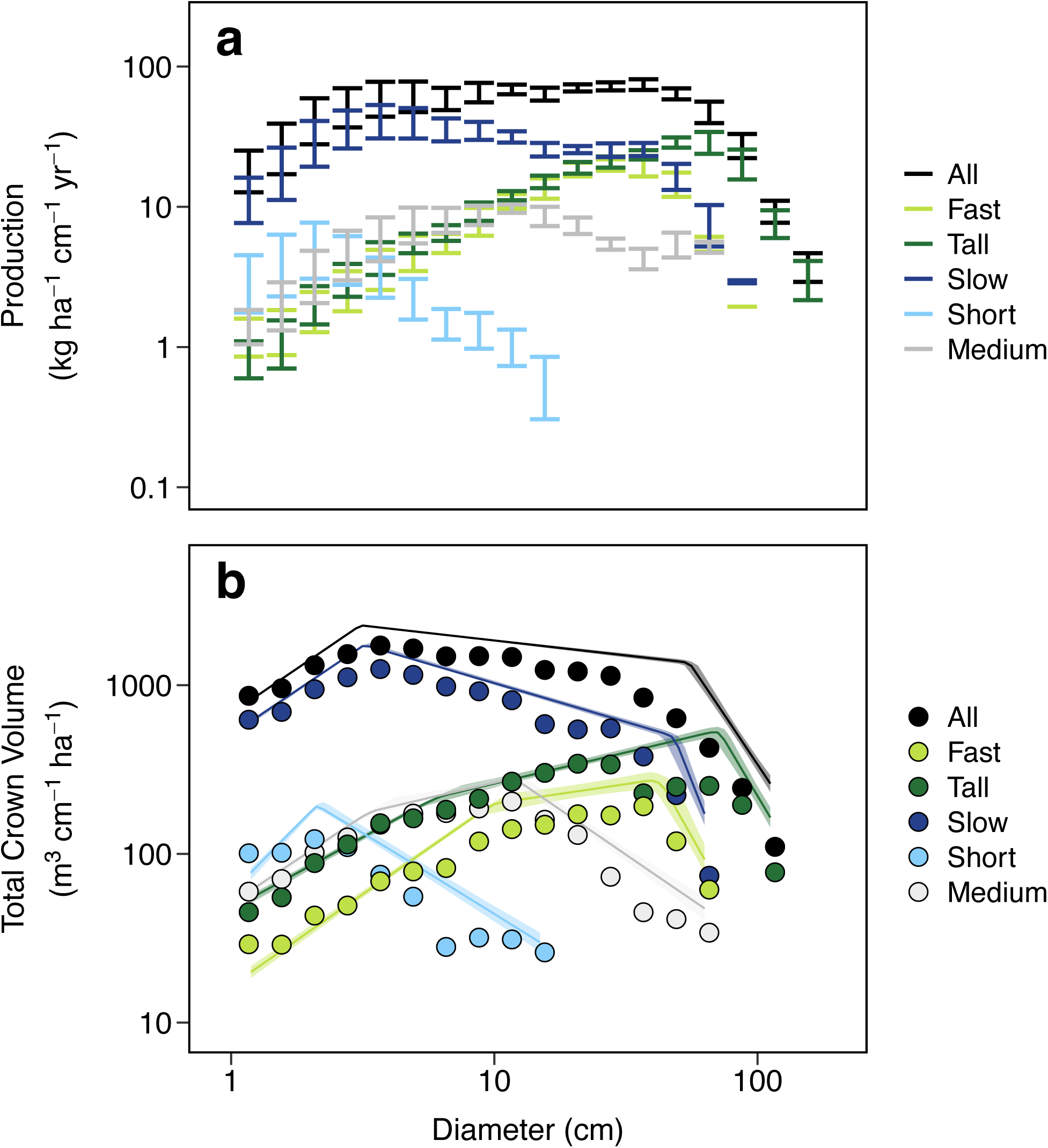
Range of Biomass Growth and Crown Volume. **a** Ranges in production from 6 censuses spanning 1985-2010. **b** Total crown volume in 1995.

**Fig. S7.**
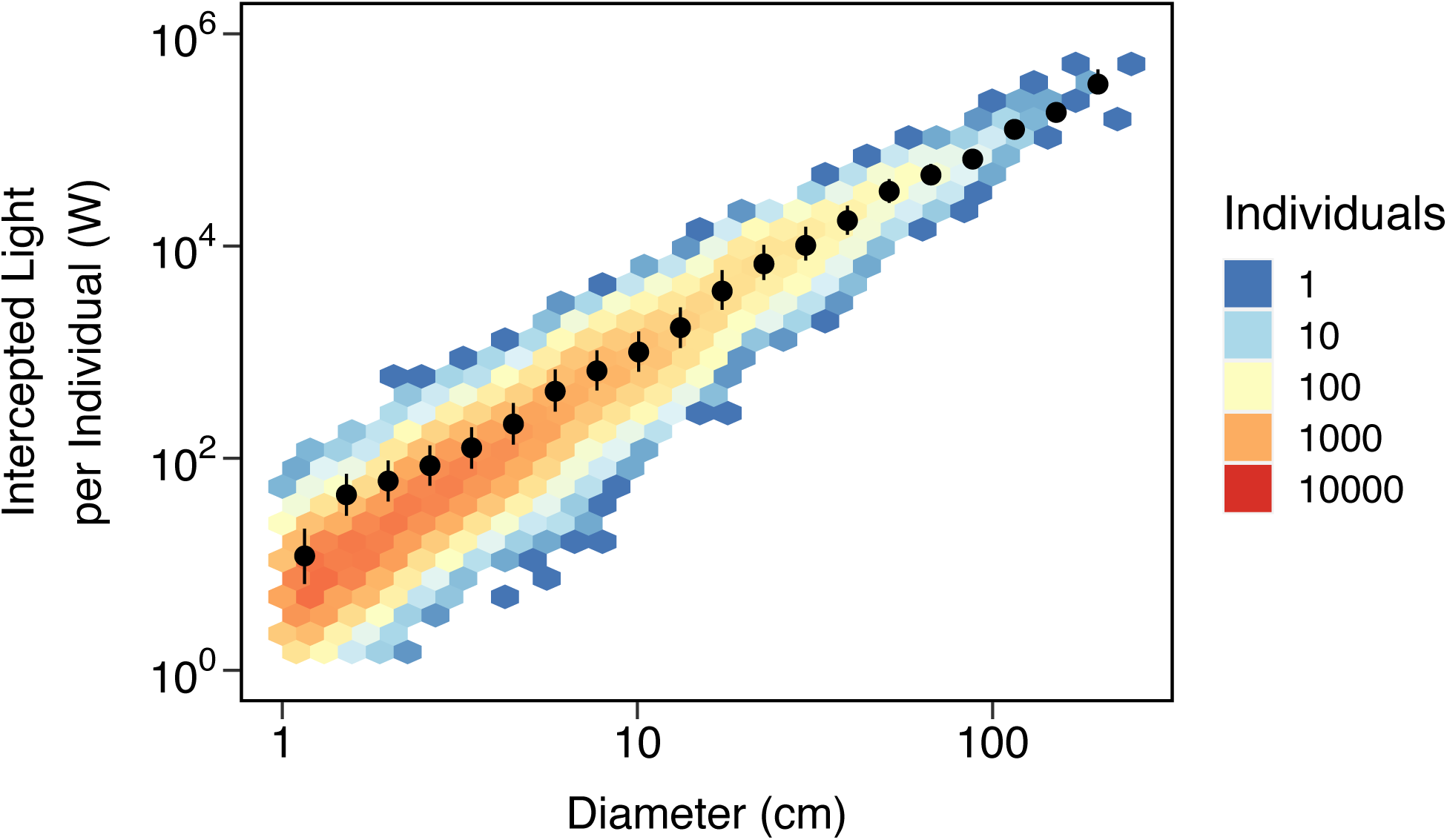
Individual light capture with size. Light capture follows a power law; mean values with 25% and 75% quantiles are shown in black. Multiplication of individual light capture regression fit with population density (Fig. 4b) yields total light capture plotted in Fig. 5a.

**Fig. S8.**
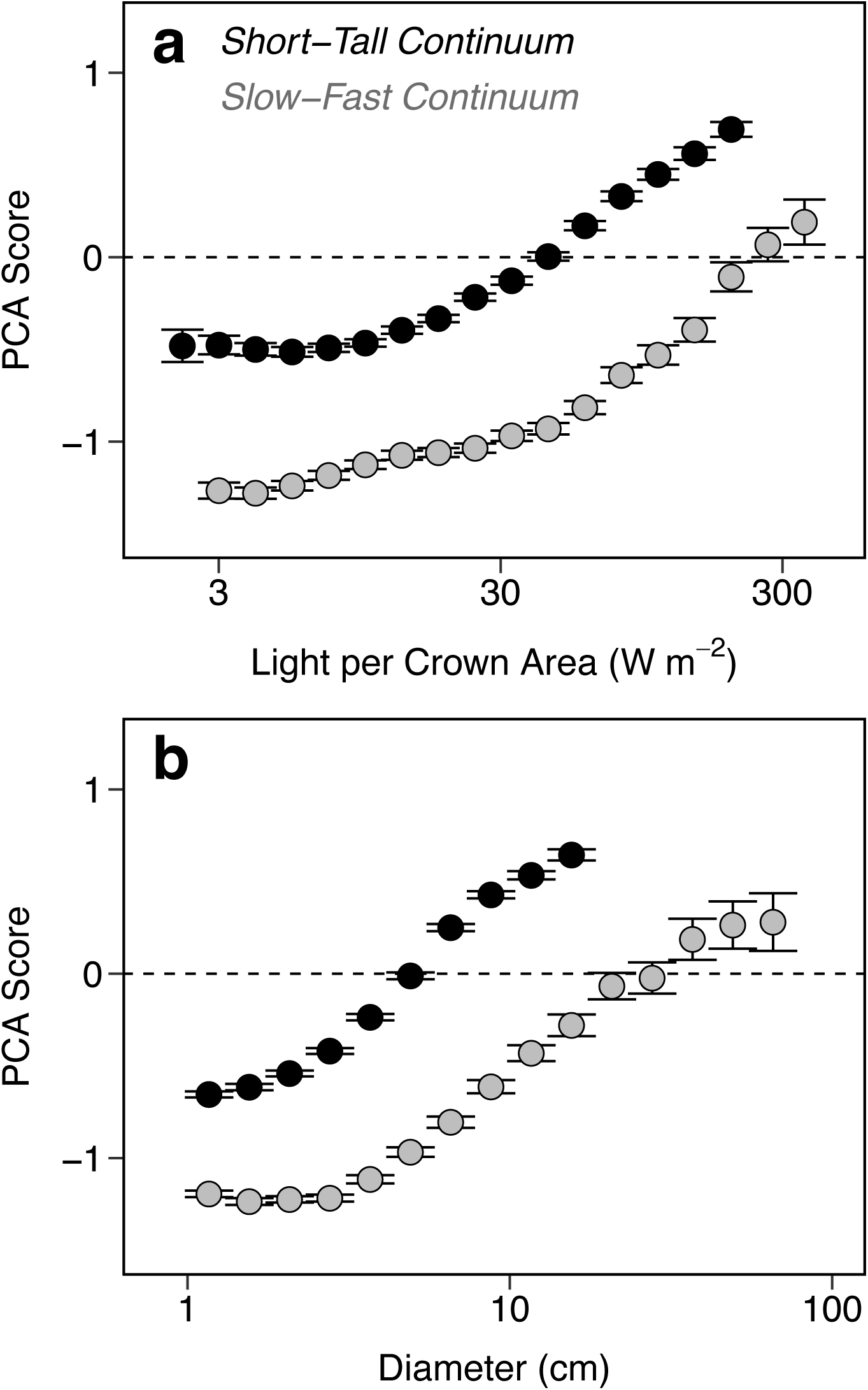
Mean life history PCA scores across size and light. **a-b** Mean PCA scores for all individuals along two life history dimensions are plotted with 95% credible intervals.

**Fig. S9.**
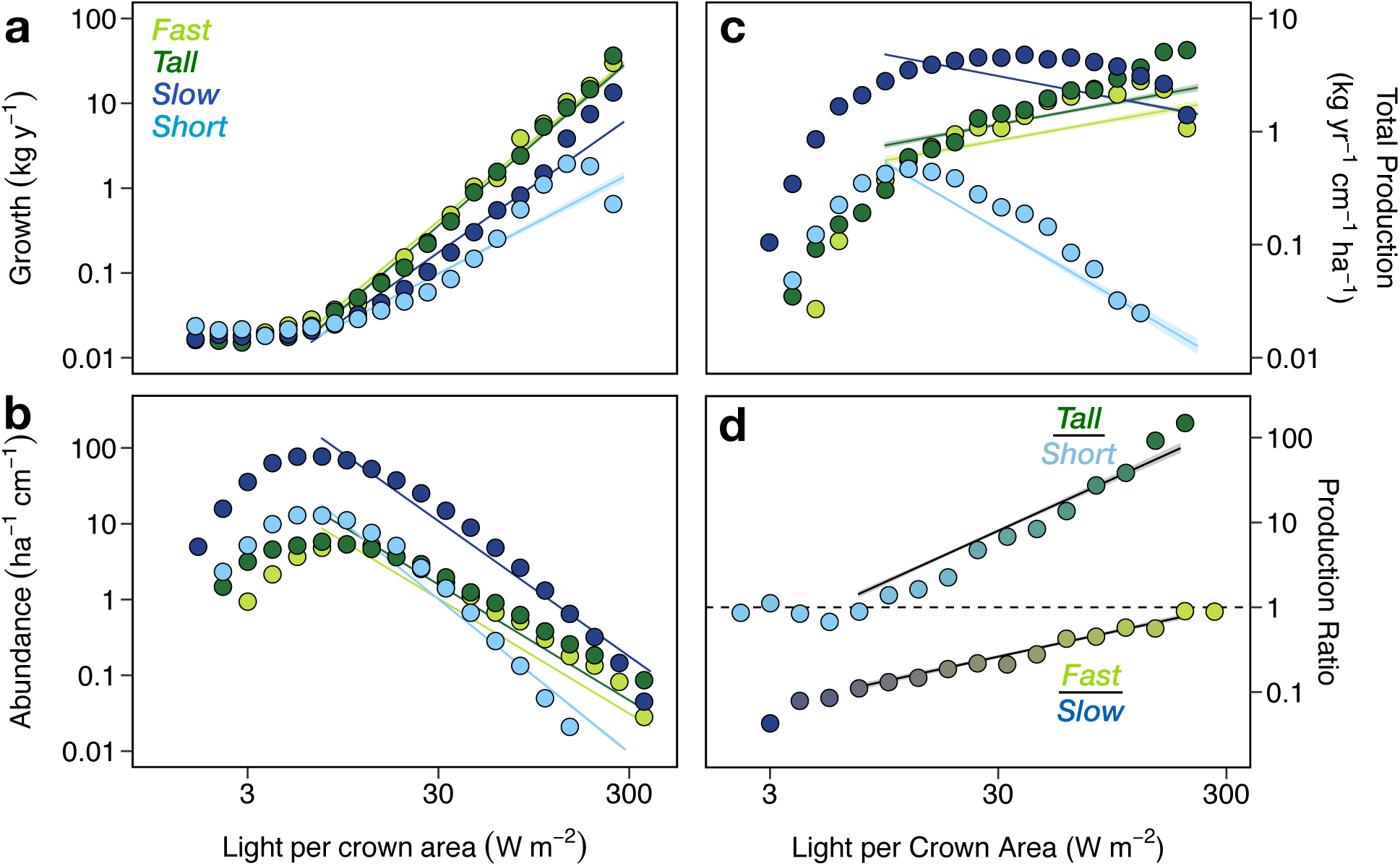
Growth, abundance and production with light. Linear regression fits for **a** growth and **b** abundance are multiplied to estimate **c** production for respective life histories. Regression fits in **c** are used to calculate dimensionless ratio fits in **c** and Fig. 6a. 95% credible bands are shown on all regressions, and regression fits are limited to > 7 W m^−2^, where patterns are approximately linear. Note that despite a two order of magnitude decline in production below 7 W m^−2^ in **c**, dimensionless life history production ratios are approximately linear across the full range of light conditions in **d**, consistent with theoretical predictions (Eq. 5).

## References

1 Gravel, D., Canham, C. D., Beaudet, M. & Messier, C. Shade tolerance, canopy gaps and mechanisms of coexistence of forest trees. Oikos 119, 475–484 (2010).

2 Farrior, C., Bohlman, S., Hubbell, S. & Pacala, S. Dominance of the suppressed: Power-law size structure in tropical forests. Science 351, 155–157 (2016).

3 Kobe, R. K., Pacala, S. W., Silander, J. A. & Canham, C. D. Juvenile tree survivorship as a component of shade tolerance. Ecological applications 5, 517–532 (1995).

4 Denslow, J. S. Gap partitioning among tropical rainforest trees. Biotropica, 47–55 (1980).

5 Kobe, R. K. Light gradient partitioning among tropical tree species through differential seedling mortality and growth. Ecology 80, 187–201 (1999).

6 Montgomery, R. A. & Chazdon, R. L. Forest structure, canopy architecture, and light transmittance in tropical wet forests. Ecology 82, 2707–2718 (2001).

7 Balderrama, S. I. V. & Chazdon, R. L. Light-dependent seedling survival and growth of four tree species in Costa Rican second-growth rain forests. Journal of Tropical Ecology 21, 383–395 (2005).

8 Montgomery, R. & Chazdon, R. Light gradient partitioning by tropical tree seedlings in the absence of canopy gaps. Oecologia 131, 165–174 (2002).

9 Kitajima, K. & Poorter, L. Functional basis for resource niche partitioning by tropical trees. Tropical forest community ecology, 160–181 (2008).

10 Bonsall, M. B., Jansen, V. A. & Hassell, M. P. Life history trade-offs assemble ecological guilds. Science 306, 111–114 (2004).

11 Ricklefs, R. E. Environmental heterogeneity and plant species diversity: a hypothesis. The American Naturalist 111, 376–381 (1977).

12 Wright, S. J. et al. Functional traits and the growth–mortality trade-off in tropical trees. Ecology 91, 3664–3674 (2010).

13 Reich, P. B. The world-wide ‘fast–slow’ plant economics spectrum: a traits manifesto. Journal of Ecology 102, 275–301 (2014).

14 Sterck, F., Markesteijn, L., Schieving, F. & Poorter, L. Functional traits determine trade-offs and niches in a tropical forest community. Proceedings of the National Academy of Sciences 108, 20627–20632 (2011).

15 Poorter, L. & Bongers, F. Leaf traits are good predictors of plant performance across 53 rain forest species. Ecology 87, 1733–1743 (2006).

16 Hubbell, S. P. The Unified Neutral Theory of Biodiversity and Biogeography. Vol. 32 (Princeton University Press, 2001).

17 Hubbell, S. P. et al. Light-gap disturbances, recruitment limitation, and tree diversity in a neotropical forest. Science 283, 554–557 (1999).

18 Lieberman, M., Lieberman, D., Peralta, R. & Hartshorn, G. S. Canopy closure and the distribution of tropical forest tree species at La Selva, Costa Rica. Journal of Tropical Ecology 11, 161–177 (1995).

19 Mangan, S. A. et al. Negative plant–soil feedback predicts tree-species relative abundance in a tropical forest. Nature 466, 752 (2010).

20 Bagchi, R. et al. Pathogens and insect herbivores drive rainforest plant diversity and composition. Nature 506, 85 (2014).

21 Forrister, D. L., Endara, M.-J., Younkin, G. C., Coley, P. D. & Kursar, T. A. Herbivores as drivers of negative density dependence in tropical forest saplings. Science 363, 1213–1216, doi: 10.1126/science.aau9460 (2019).

22 Uhl, C., Clark, K., Dezzeo, N. & Maquirino, P. Vegetation dynamics in Amazonian treefall gaps. Ecology 69, 751–763 (1988).

23 Muller-Landau, H. et al. Comparing tropical forest tree size distributions with the predictions of metabolic ecology and equilibrium models. Ecology Letters 9, 589–602 (2006).

24 Perkins, D. M. et al. Energetic equivalence underpins the size structure of tree and phytoplankton communities. Nature Communications 10, 255 (2019).

25 Enquist, B., West, G., Charnov, E. & Brown, J. Allometric scaling of production and life-history variation in vascular plants. Nature 401, 907–911 (1999).

26 Hatton, I. A., Dobson, A. P., Storch, D., Galbraith, E. D. & Loreau, M. Linking scaling laws across eukaryotes. Proceedings of the National Academy of Sciences 116, 21616–21622 (2019).

27 Zhang, W.-P., Morris, E. C., Jia, X., Pan, S. & Wang, G.-X. Testing predictions of the energetic equivalence rule in forest communities. Basic and Applied Ecology 16, 469–479 (2015).

28 Belgrano, A., Allen, A. P., Enquist, B. J. & Gillooly, J. F. Allometric scaling of maximum population density: a common rule for marine phytoplankton and terrestrial plants. Ecology letters 5, 611–613 (2002).

29 West, G. B., Enquist, B. J. & Brown, J. H. A general quantitative theory of forest structure and dynamics. Proceedings of the National Academy of Sciences 106, 7040–7045 (2009).

30 Taubert, F., Jahn, M. W., Dobner, H.-J., Wiegand, T. & Huth, A. The structure of tropical forests and sphere packings. Proceedings of the National Academy of Sciences 112, 15125–15129 (2015).

31 Coomes, D. A., Lines, E. R. & Allen, R. B. Moving on from Metabolic Scaling Theory: hierarchical models of tree growth and asymmetric competition for light. Journal of Ecology 99, 748–756 (2011).

32 Stark, S. C. et al. Linking canopy leaf area and light environments with tree size distributions to explain Amazon forest demography. Ecology letters 18, 636–645 (2015).

33 Grady, J. M. et al. Metabolic asymmetry and the global diversity of marine predators. Science 363, eaat4220 (2019).

34 Condit, R. et al. Barro Colorado Forest Census Plot Data, 2012 Version. DataONE Dash, doi: http://dx.doi.org/10.5479/data.bci.20130603 (2012).

35 Rüger, N. et al. Beyond the fast–slow continuum: demographic dimensions structuring a tropical tree community. Ecology Letters 21, 1075–1084, doi: 10.1111/ele.12974 (2018).

36 Rüger, N., Huth, A., Hubbell, S. P. & Condit, R. Determinants of mortality across a tropical lowland rainforest community. Oikos 120, 1047–1056 (2011).

37 Kitajima, K., Mulkey, S. S. & Wright, S. J. Variation in crown light utilization characteristics among tropical canopy trees. Annals of Botany 95, 535–547 (2005).

38 Valladares, F. & Niinemets, Ü. Shade tolerance, a key plant feature of complex nature and consequences. Annual Review of Ecology, Evolution, and Systematics 39 (2008).

39 Hurlbert, A. H. Species–energy relationships and habitat complexity in bird communities. Ecology Letters 7, 714–720 (2004).

40 White, E. P., Enquist, B. J. & Green, J. L. On estimating the exponent of power-law frequency distributions. Ecology 89, 905–912 (2008).

41 Bazzaz, F. The physiological ecology of plant succession. Annual review of Ecology and Systematics 10, 351–371 (1979).

42 Hubbell, S. P. & Foster, R. B. Short-term dynamics of a neotropical forest: why ecological research matters to tropical conservation and management. Oikos, 48–61 (1992).

43 Rüger, N. et al. Demographic trade-offs predict tropical forest dynamics. Science 368, 165–168 (2020).

44 Brown, J. H., Gillooly, J. F., Allen, A. P., Savage, V. M. & West, G. B. Toward a metabolic theory of ecology. Ecology 85, 1771–1789 (2004).

45 Enquist, B. J. et al. Scaling metabolism from organisms to ecosystems. Nature 423, 639–642 (2003).

46 Tilman, D. et al. Does metabolic theory apply to community ecology? It’s a matter of scale. Ecology 85, 1797–1799 (2004).

47 White, E. P., Ernest, S., Kerkhoff, A. J. & Enquist, B. J. Relationships between body size and abundance in ecology. Trends in Ecology & Evolution 22, 323–330 (2007).

48 Grady. Methods for ‘Life history scaling and the division of energy in forests’. (2020).

49 Vermeij, G. J. Inequality and the directionality of history. The American Naturalist 153, 243–253 (1999).

50 Levine, J. M., Bascompte, J., Adler, P. B. & Allesina, S. Beyond pairwise mechanisms of species coexistence in complex communities. Nature 546, 56–64 (2017).

51 Windsor, D. M. in Ecología de un bosque tropical: ciclos estacionales y cambios a largo plazo (ed A. S. Rand) 53–71 (Smithsonian Trop. Res. Inst.,, 1990).

52 Condit, R. Tropical forest census plots: methods and results from Barro Colorado Island, Panama and a comparison with other plots. (Springer Science & Business Media, 1998).

53 Foster, R. & Brokaw, N. in The Ecology of a Tropical Forest: Seasonal Rhythms and Long-Term Changes. (eds EG Leigh Jr., AS Rand, & DM Windsor) 67–81 (1982).

54 Meakem, V. et al. Role of tree size in moist tropical forest carbon cycling and water deficit responses. New Phytologist (2017).

55 Cushman, K., Muller-Landau, H. C., Condit, R. S. & Hubbell, S. P. Improving estimates of biomass change in buttressed trees using tree taper models. Methods in Ecology and Evolution 5, 573–582 (2014).

56 Metcalf, C. J. E., Clark, J. S. & Clark, D. A. Tree growth inference and prediction when the point of measurement changes: modelling around buttresses in tropical forests. Journal of Tropical Ecology 25, 1–12 (2009).

57 Feldpausch, T. R. et al. Tree height integrated into pantropical forest biomass estimates. Biogeosciences, 3381–3403 (2012).

58 Cano, I. M., Muller-Landau, H. C., Wright, S. J., Bohlman, S. A. & Pacala, S. W. Tropical tree height and crown allometries for the Barro Colorado Nature Monument, Panama: a comparison of alternative hierarchical models incorporating interspecific variation in relation to life history traits. Biogeosciences 16, 847–862 (2019).

59 Bohlman, S. & O’Brien, S. Allometry, adult stature and regeneration requirement of 65 tree species on Barro Colorado Island, Panama. Journal of Tropical Ecology 22, 123–136 (2006).

60 Sprugel, D. Correcting for bias in log-transformed allometric equations. Ecology 64, 209–210 (1983).

61 Chave, J. et al. Tree allometry and improved estimation of carbon stocks and balance in tropical forests. Oecologia 145, 87–99 (2005).

62 Wirth, R., Weber, B. & Ryel, R. J. Spatial and temporal variability of canopy structure in a tropical moist forest. Acta Oecologica 22, 235–244 (2001).

63 North, G. R. Analytical solution to a simple climate model with diffusive heat transport. Journal of the Atmospheric Sciences 32, 1301–1307 (1975).

64 Devadoss, S., Luckstead, J., Danforth, D. & Akhundjanov, S. The power law distribution for lower tail cities in India. Physica A: Statistical Mechanics and its Applications 442, 193–196 (2016).

65 Gelman, A., Goodrich, B., Gabry, J. & Vehtari, A. R-squared for Bayesian regression models. The American Statistician, 1–7 (2019).

66 Denny, M. The fallacy of the average: on the ubiquity, utility and continuing novelty of Jensen’s inequality. Journal of Experimental Biology 220, 139–146 (2017).

67 Grady, J. M., Enquist, B. J., Dettweiler-Robinson, E., Wright, N. A. & Smith, F. A. Evidence for mesothermy in dinosaurs. Science 344, 1268–1272 (2014).

68 Gillooly, J., Charnov, E., West, G., Savage, V. & Brown, J. Effects of size and temperature on developmental time. Nature 417, 70–73 (2002).

69 Hamilton, M. J., Davidson, A. D., Sibly, R. M. & Brown, J. H. Universal scaling of production rates across mammalian lineages. Proceedings of the Royal Society B: Biological Sciences 278, 560–566 (2011).

70 Meiri, S., Brown, J. H. & Sibly, R. M. The ecology of lizard reproductive output. Global Ecology and Biogeography 21, 592–602 (2012).

71 Sterck, F., Poorter, L. & Schieving, F. Leaf traits determine the growth-survival trade-off across rain forest tree species. The American Naturalist 167, 758–765 (2006).

72 Team, S. D. RStan: the R interface to Stan. R package version 2.18.2. http://mc-stan.org/. (2018).

73 Vehtari A, Gabry J, Yao Y & Gelman A loo: Efficient leave-one-out cross-validation and WAIC for Bayesian models. R package version 2.1.0, https://CRAN.R-project.org/package=loo (2019).

74 Gelman, A. & Rubin, D. B. Inference from iterative simulation using multiple sequences (with discussion). Statistical Science 7, 457–511 (1992).

